# REST elevation-dependent chromatin remodeling and alternative Grk6 transcript synthesis hyperactivates Cxcr4-Sdf1 signaling in cerebellar granule cell progenitors

**DOI:** 10.1101/2025.10.15.682654

**Authors:** Keri Callegari, Jyothishmathi Swaminathan, Lei Guo, Ashutosh Singh, Yanwen Yang, Xue Xiao, Tara H.W. Dobson, Amanda R. Haltom, Javiera Bravo-Alegria, Ajay Sharma, Xin Hu, Lin Xu, Vidya Gopalakrishnan

## Abstract

*RE1* Silencing Transcription Factor (REST) is a repressor of transcriptional initiation of genes involved in neurogenesis. Here, we show that conditional *REST* elevation in cerebellar granule cell progenitors (CGNPs) of *REST^TG^* mice perturbed foliation, increased cell migration, and sustained C-X-C motif receptor 4 (Cxcr4) signaling, a pathway key to postnatal CGNP migration. Mechanistic studies uncovered a novel role for REST in controlling transcript diversity and exon skipping in CGNPs. Alternative transcript expression was detected in known Cxcr4 signaling regulator, G-protein-coupled receptor kinase-6 (*Grk6*). Further analysis of Grk6’s isoform expression revealed an upregulation of a transcript lacking exon 10a (*Grk6-207*) in *REST^TG^* CGNPs. *Grk6-207* expression in wildtype CGNPs hyperactivated Cxcr4 signaling and increased chemotaxis. Structural modeling of Grk6-207 predicted changes in active site conformation and interactions with Cxcr4 and β-Arrestin-1, supporting impairment of Cxcr4 signaling desensitization. Interestingly, *REST* elevation promoted increased chromatin accessibility at the exon10a-10b junction and exon 10a exclusion. Integrated multiomic analyses identified the enhancer of zeste (Ezh2) as a potential mediator of alternative transcript generation which demonstrated increased occupancy at the exon10a-10b locus in *REST^TG^* CGNPs. Pharmacological inhibition of Ezh2 downregulated *Grk6-207*, confirming a role for Ezh2 in *Grk6* exon10a exclusion and the increased migration in *REST^TG^* CGNPs.

**GRAPHICAL ABSTRACT:** 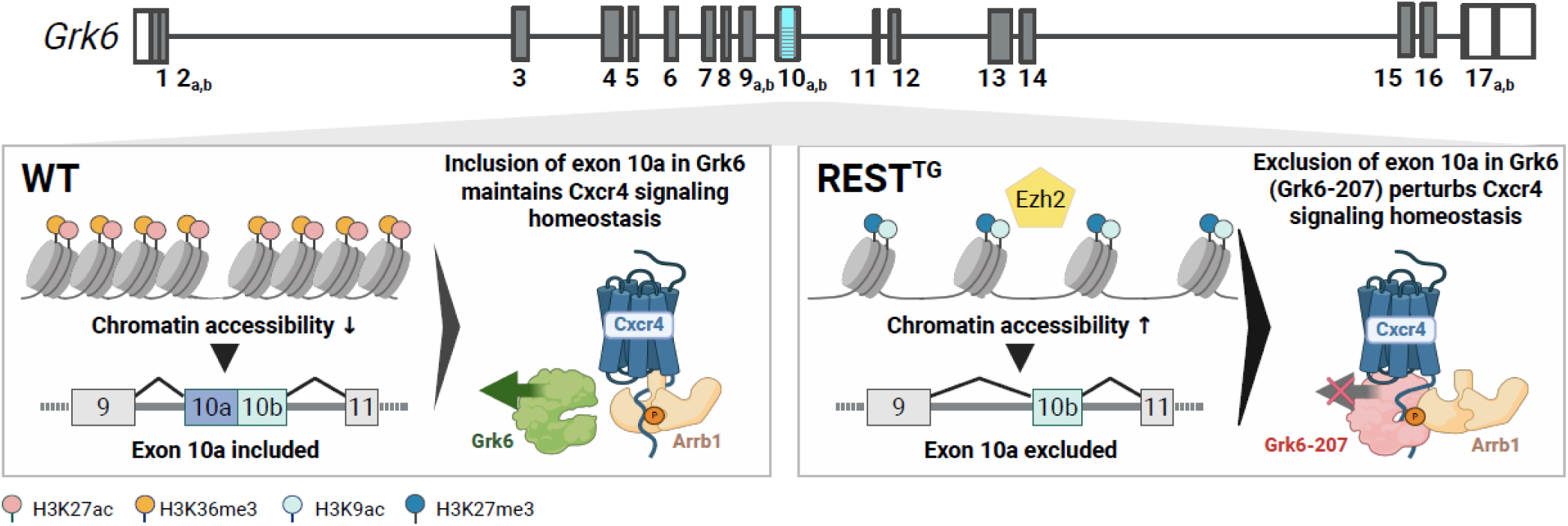

## INTRODUCTION

The *RE1* Silencing Transcription Factor (REST) is a canonical transcriptional repressor of neuronal differentiation genes (1,2). REST plays critical roles in neurodevelopment and neurogenesis and both under- and over-expression of REST is associated with pathological phenotypes (3-6). Abnormal *REST* levels and/or nuclear localization is noted in Huntington’s disease, neurodevelopmental conditions like autism, and pediatric brain cancers (neuroblastoma, glioblastoma, and subsets of medulloblastoma (MB)) (7-12). We and others previously showed that elevated REST in SHH subgroup of MBs is associated with metastasis and leptomeningeal dissemination, which are major causes of poor patient survival (13-15). However, the underlying mechanisms are still under investigation.

Here, we studied the effects of REST elevation in cerebellar granule neuron progenitors (CGNPs), the cells of origin of SHH-MBs, using a previously described REST transgenic mouse model (*REST^TG^*) (16). A detailed characterization of the mouse model revealed defects in cerebellar foliation and lamination, associated with perturbed localization of CGNPs. The above phenotype was associated with increased expression of the chemokine receptor C-X-C motif receptor 4 (Cxcr4) expression and activity in response to its ligand stromal-cell derived factor 1 (Sdf1), both of which are implicated in cerebellar development and MB genesis (17-20). During postnatal cerebellar development, Cxcr4-Sdf1 signaling restricts CGNPs to the proliferative external granule cell layer (EGL) and promotes their tangential migration (21). Targeted deletion of Cxcr4 results in severe disorganization of the cerebellum while Cxcr4 overactivation results in foliation defects which are also observed in human warts hypogammaglobulinemia immunodeficiency myelokathexis (WHIM) syndrome patients (18). Further, deregulation and hyperactivation of CXCR4 is correlated with metastasis in SHH-MB (22-25).

REST/Rest is a known repressor of transcriptional initiation and acts in concert with two distinct repressive chromatin remodeling complexes - mSin3a and co-REST, bound to its amino (-N) and carboxy (-C) termini, respectively. REST-mediated gene silencing involves the activity of enzymes such as histone deacetylases, histone H3 lysine (K)-9 methyltransferases (G9a and G9a-like protein), histone H3 K4 demethylases (LSD1) and the SWI/SNF ATP-dependent chromatin remodelers found within the mSIN3A and co-REST complexes (26-29). While most functional studies have examined REST’s repressive activity at target gene promoters, the functional relevance of REST activity at intra- and intergenic locations is underexplored (30-32).

In this study, we used multiomic analyses and functional validation in driving the synthesis of alternate isoforms of genes involved in controlling CGNPs migration in *REST^TG^* mice. Specifically, we describe how REST-dependent generation of an alternative transcript for the gene encoding the G-protein coupled receptor kinase 6 (*Grk6*) called *Grk6*-207, which lacks exon 10a, results in sustained Cxcr4 phosphorylation and pathway hyperactivity (25,33,34). Structural modeling studies suggest that the increased affinity of the protein encoded by *Grk6-207* transcript may reduce Cxcr4 phosphorylation-dependent recruitment of the chaperone - beta-arrestin (Arrb1) and signaling desensitization to drive constitutively high activity of Cxcr4. We also demonstrate a role for chromatin remodeling in the REST-elevation dependent generation of *Grk6-207.* In particular, REST elevation in CGNPs was associated with an unexpected increase in accessibility of chromatin 5’ proximal to exon 10a-exon10b of *Grk6*, aberrant occupancy of this locus by the Enhancer of Zeste (Ezh2), deposition of histone H3K27 trimethylation (me3) and a reduction in histone H3K36me3 – a mark of active transcription. The reversal of these changes by the EZH2 inhibitor, Tazemetostat, suggest a role for Ezh2 in REST-mediated aberrant *Grk6-207* expression, sustained Cxcr4-Sdf1 signaling and cell migration in CGNPs in mice.

## MATERIALS AND METHODS

### Animals

*REST^TG^* mice were generated as described previously. *hREST* transgene was induced by intraperitoneal (IP) injection of tamoxifen on postnatal (P) days 2, 3 and 4 (16). Mice were housed and treated in accordance with The University of Texas MD Anderson Cancer Center’s Institutional Animal Care and Use Committee (IACUC) guidelines.

### Immunohistochemistry (IHC)

Antibodies used for IHC are listed in supplemental Table S1. Secondary rabbit and mouse antibodies in glycerol were obtained from Jackson Labs (Bar Harbor ME, USA) and used at 1:50 and 1:100, respectively. Slides were counterstained with hematoxylin.

### Cell Culture & Transwell Migration Assay

Isolation and cultivation of primary CGNPs from WT and *REST^TG^* mice was performed as described previously (16). Transwell assays (BD biosciences – BioCoat Matrigel Invasion Chambers – 354480) were performed using CGNPs with and without the presence of recombinant SDF1 (rSDF1) ligand (100ng/mL) and with and without AMD3100 (2.6μM) for 24 hours.

### Cloning of *Grk6* isoforms

Gateway cloning method was used to create the *mGrk6* lentivirus expression vectors. *mGrk6-202* was assembled from synthetic oligonucleotides and/or PCR products and inserted into the phage- *ef1a*-m*Grk6*-202-IRES-*eGFP* (or *mkate2*) lentivirus expression vector. For generation of the m*Grk6*- 207 exon 10 fragment, we performed overlap PCR to deleted 102bp nt from the m*Grk6*-202. Control lentivirus expression vector phage-*ef1a*- Flag-IRES-*eGFP* (or *mkate2*) was created by cloning the Flag insert alone into the vector.

The Lentivirus packaging 293T/17 cells (ATCC® CRL-11268™) cells were transfected with phage- *ef1a-*m*GRK6-202*-IRES-*eGFP* (or *mkate2*) or phage-*ef1a-* m*GRK6-207*-IRES-*eGFP* (or *mkate2*) or phage-*ef1a*- Flag-IRES-*eGFP* (or *mkate2*) with packaging vector *psPAX2* (Addgene plasmid # 12260) and *pVSV-G* (Clontech) by using Lipofectamine 2000, according to manufacturer’s instructions. Ex vivo expanded mouse CGNPs were seeded in 6 well plates in 1.5ml progenitor cell proliferation media. For transduction, 0.5ml viral supernatants were added to CGNP cultures at a final volume of 2ml and incubated in 34°C for 24 hours. Transduction efficiency was verified by fluorescence microscopy imaging. m*Grk6-202*, m*Grk6-207* expression was confirmed using qPCR.

### Western Blotting

Western blotting was performed using ex vivo expanded CGNPs harvested from WT and *REST^TG^* mice as described previously (16). Primary antibodies used for analyses are listed in supplemental Table S2.

### qRT-PCR

Equal amounts of RNA, up to 1 μg, isolated using Zymo Research Quick-RNA mini-prep kit (Zymo Research – R1055), were reverse-transcribed into cDNA using the iScript cDNA Synthesis Kit (Bio-Rad, Hercules, CA). Quantitative RT-PCR was performed in triplicate with a 2X SensiMix SYBR &

Fluorescein Kit (Bioline, Boston, MA) using a LightCycler 96 Real-Time PCR System (Roche Diagnostics GmbH, Mannheim, Germany). Relative mRNA expression normalized to 18S ribosomal RNA was determined by the comparative 2-ΔΔCp method and graphed as fold change compared to WT controls (16). Primers sequences are listed in supplemental Table S3.

### RNAseq analysis

RNA was harvested from CGNPs pooled from WT and *REST^TG^* mice (n=10 each) as above and quality checked using RNA Nano assay using Bioanalyzer 2100 (Agilent Technologies - RNA RIN should be >8). Quantification was done using Qubit 2.0 Fluorometer (Thermo Fisher Scientific, Waltham, MA). Library preparation was done with 1000 ng of RNA using TruSeq Stranded Total RNA Gold kit (Illumina RS-122-2303) following TruSeq Stranded Total RNA Reference guidelines. RNA-seq library quality/quantity was checked using Tape Station 2200 D1000 assay (Agilent Technologies, Santa Clara, CA and QuantStudio 6 Flex Real time PCR System (Applied Biosystem, Waltham, MA), and sequenced (paired end-75 cycles) using Hiseq 3000 (Illumina, San Diego, CA). Adapters were trimmed using Trim Galore (v0.6.4). Isoform quantification was performed with Salmon (v1.10.1). Isoforms with zero expression in any sample were excluded from further analysis. Expression data were normalized using the Voom method from the R package limma (v3.50.3). Differentially expressed isoforms were identified using limma, and Gene Set Enrichment Analysis (GSEA) for differentially expressed genes (DEGs) was conducted using the R package clusterProfiler (v4.2.2). For downstream analysis, genes with alternative isoform usage were identified by evaluating significant expression differences between wildtype and *REST^TG^* conditions and prioritizing genes which exhibited changes in non-APPRIS transcripts. Individual analyses of exonic (FPKM) and junction coverage (MISO) were completed for prioritized genes to examine alterations in specific transcript usage, which overcomes the limitations of probabilistic mapping of reads to alternative transcripts contained within full-length transcripts.

For junction analysis and generation of Sashimi plots to obtain quantitative estimates of isoform abundance, reads were aligned to the reference genome mm10 in the standard BAM format. Splice junction alignments were produced using Tophat2 and alternatively spliced events annotated using MISO (35). Sashimi plots were then generated to show differential *Grk6* transcript abundance between WT and REST^TG^ CGNPs (36).

### Structural Modeling with AlphaFold3

*In silico* structural biology experiments were performed using AlphaFold3, an AI-powered computational approach which predicts tertiary and quaternary protein structures using primary sequences (Jumper et al., 2021; Veradi et al., 2021). Primary sequences were obtained from NCBI and Uniprot for Cxcr4 (P70658), Grk6-202 (Q9EP84), Grk6-207 (A0A286YE10), and beta-arrestin (Arrb1; J3QNU6) and predictions were made with the presence of phosphorylated serine 346 (S346-p), ATP, and ADP.

### ATACseq

ATAC-seq was performed in duplicate using the Omni-ATAC protocol as described (37). Briefly, nuclei were isolated from 50,000 WT and *REST^TG^* CGNPs collected in lysis buffer containing 0.1% NP-40 and 0.01% digitonin, and tagmented with Nextera Tn5 Transposase (TDE1, Illumina) in TD Tagment DNA buffer (Illumina). Resulting library fragments were purified using a Qiagen MinElute kit (Qiagen, Hilden, Germany), amplified for 4–6 PCR cycles (37), purified using AMPure XP beads (Beckman Coulter, Brea, CA) and sequenced 2 x 75 bp on an Illumina HiSeq3000 to obtain at least 50 million high quality mapping reads per sample. For mouse ATACseq datasets, paired-end raw reads were mapped to the mouse mm10 reference dataset, using bowtie2 (version 2.3.4.3) with parameter ‘–very-sensitive’ enabled (38). Read duplication and reads that mapped to chrM were removed from downstream analysis. Peaks were called using findpeaks command from HOMER software package version 4.9, with parameter ‘–style dnase’, and the FDR threshold (for poisson p-value cutoff) was set to 0.00128. Called peaks were merged from all samples and annotatePeaks.pl command was used to produce raw count matrix. Differential peaks were identified using R package DEseq version 3.82. Peaks with > 2-fold change were designated as DE peaks. To analyze the functional significance of peaks, Genomic Regions Enrichment of Annotations Tool (GREAT) (39). was used with mm10 as the background genome and other parameters set as default for mouse.

### Integration of RNAseq and ATACseq data

A previously published Bayesian-based statistical framework which integrates multiple types of cancer genomic data was adapted to perform an integrated analysis of RNAseq and ATACseq data (40,41). The approach has the advantage of avoiding false positives and negatives brought in by more traditional linear correlation and overestimation of false discovery rate (FDR) by permutation-based tests (40,42-44). In this Bayesian-based pipeline, expression of each gene was divided into groups without or with open chromatin, separately. Group differences in gene expression were then compared by Bayesian statistics instead of permutation-based test (or other frequentist statistics). The *Markov chain Monte Carlo* (MCMC) method was then used to calculate the posterior distribution, *p*(*θ*|*y*), where *p*(*θ)* is the prior probability and *p*(*y)* is the marginal likelihood. The highest density interval (*I*) was calculated based on the following equation,

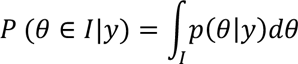

which provides posterior probability estimation of group differences in gene expression. The statistical results and other information were then extracted and summarized from Bayesian estimation. A stringent criterion (probability that group difference is greater than zero must be more than 99.9% in Bayesian estimation) was used to define statistically significant association between gene expression and open chromatin status (40).

### Transcription factor (TF) motif analysis

ChIP-seq peak datasets of 616 mouse TFs, including the ones shown in this study (REST, EZH2, SUZ12, c-MYC, E2F1, MEIS1 and FLI1), were collected from the ENCODE database (45) and Cistrome database (46). The raw reads were aligned to the mouse reference genome (GRCh38/mm10) using default parameters in BWA v0.7.12 (47). The aligned reads were subsequently filtered for quality and uniquely mappable reads were retained for further analysis using Samtools version 1.3 (48). Library complexity was measured using BEDTools version 2.26.0 (39) and meets ENCODE data quality standards (38). Peaks were called using findpeaks command from HOMER software package version 4.7 (49), similar to a previous study (50), which were used to identify target genes nearby these peaks.

### Chromatin Immunoprecipitation sequencing (ChIPseq)

Proliferating CGNPs (2-5 x10^5^ cells) from wildtype (WT) and *REST^TG^* mice (n=5, each) (for histone modifications) and wildtype (WT) and *REST^TG^* mice (n=8, each) (for transcription factor) were fixed with 1% formaldehyde, cross-linked, and processed for ChIP analyses as described previously (16).

Immunoprecipitated DNA samples were quantified by SYBRGreen qPCR using Roche Lightcycler 96 and analyzed using the comparative 2−ΔΔCp method (16). The ChIP DNA was used to prepare DNA libraries using DNA library prep kit from Cell signaling technology (Cell Signaling Technology, Danvers, CO, USA - Catalog number 56795). Briefly, about 5-50ng of ChIP DNA was end prepped using an enzyme mix and linked to an adaptor. After clean-up, PCR was used to enrich for the adaptor ligated DNA using multiplex oligos for Illumina sequencing platform (Dual index Oligos, Catalog number 47538). These concentration of DNA libraries were measured using Qubit and size distribution were determined using an Agilent Bioanalyzer High Sensitivity DNA chip, according to the manufacturer’s instructions. The braries were then pooled at equal concention of 10nM and sequenced using Illumina HiSeq3000 to obtain at least 50 million high quality mapping reads per sample. Adapters were trimmed using Trim Galore (v0.6.4). Trimmed reads were aligned to the mouse genome (mm10) using Bowtie2 (v2.4.2). Blacklisted regions in the mm10 genome were removed with Bedtools (v2.29.2). Peak calling, motif analysis, and bigwig file generation were performed using HOMER (v5.1).

### Chromatin Immunoprecipitation (ChIP)

Proliferating CGNPs (2-5 x10^5^ cells) from wildtype (WT) and *REST^TG^* mice (n=5, each) (for histone modifications) and (WT) and *REST^TG^* mice (n=8, each) (for transcription factor) were fixed with 1% formaldehyde, cross-linked, and processed for ChIP analyses as described previously (16). Immunoprecipitated DNA samples were quantified by SYBRGreen qPCR using Roche Lightcycler 96 and analyzed using the comparative 2−ΔΔCp method (16). The list of antibodies and primers are listed in the supplemental Tables S4 and S5.

## RESULTS

### REST elevation in CGNPs disrupts the cytoarchitecture of the postnatal cerebellum

To evaluate the consequences of REST elevation in CGNPs on postnatal cerebellar development, human REST transgene (hREST) expression was induced in REST^TG^ mice by tamoxifen injection on postnatal (P) days 3 – 5 (16). Brains were harvested on day P14 and coronal sections were stained with hematoxylin and eosin (H&E) to reveal measurable cerebellar asymmetry in 3 of 5 brains from *REST^TG^* mice (n=5) relative to wild type (WT) (n=3) littermates (Fig. 1A, S1A). Evaluation of sagittal sections of cerebella from *REST^TG^* mice also identified several areas of perturbed cerebellar cytoarchitecture, lamination defects, and EGL expansion compared to age matched WT controls (Fig. 1B, S1B). IHC staining of sagittal brain sections of *REST^TG^* mice revealed that most cells in the expanded EGL and some of the cells found ectopically in the leptomeningeal space were positive for REST and Pax-6 identifying these mis-localized cells as CGNPs, although cells lacking these markers were also found to be interspersed with CGNPs in these niches (Fig. 1C, S1C) (51). Further, a number of these cells were negative for the post-mitotic neuronal markers - NeuN, and Tubb3 (Fig. 1C, S1C).

**Figure 1.**
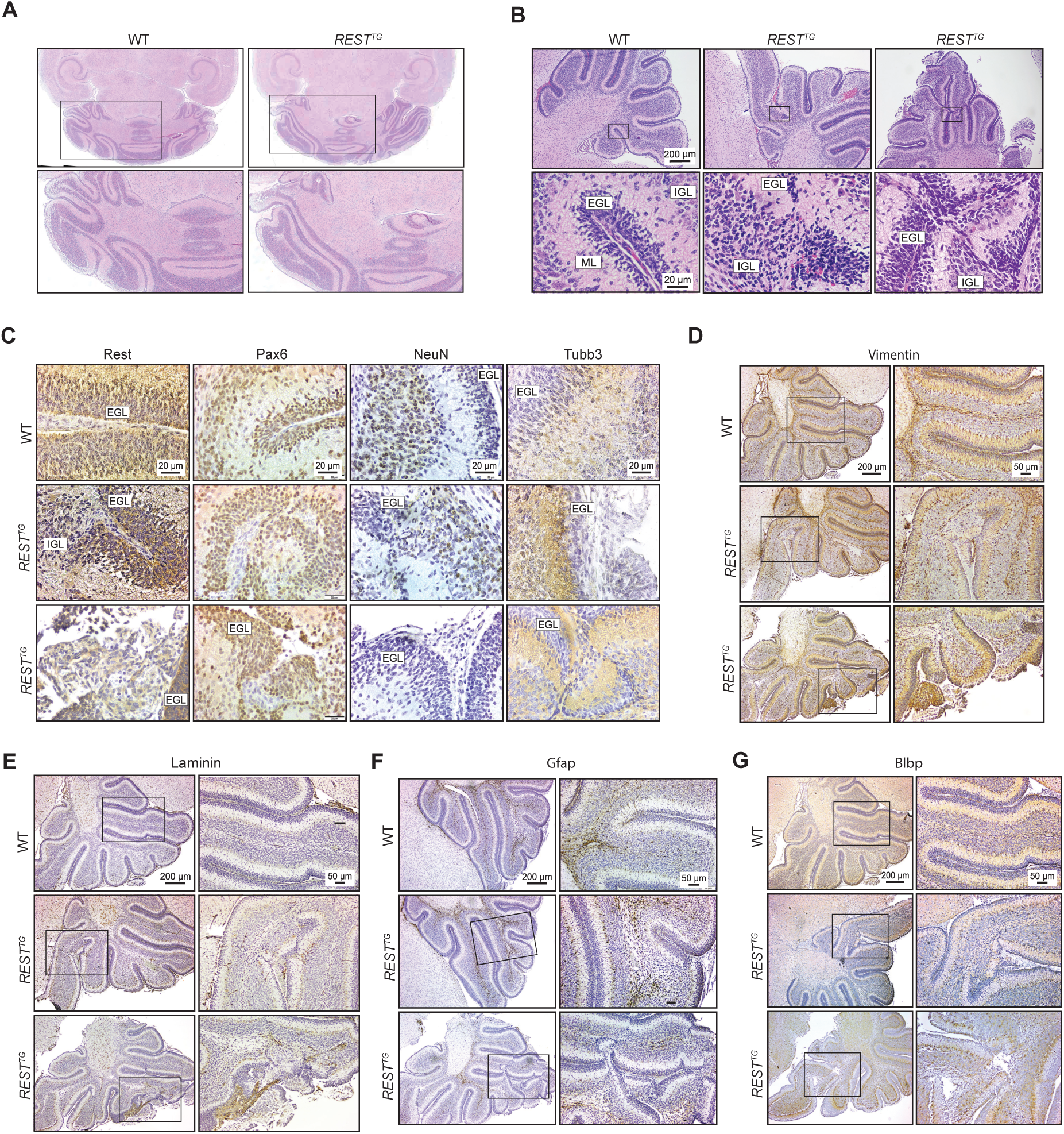
(A) Representative horizontal sections stained with hematoxylin and eosin (H&E) from WT and REST^TG^ mice (n=5, n=5; other specimens in S1A). (B) Sagittal sections stained with H&E from P8 WT (n=3) and REST^TG^ (n=3) mice. EGL=External granular layer, IGL=Internal granular cell layer, ML=Molecular layer. (C) REST, Pax-6, NeuN, Tubb3 staining, of P8 WT (n=3) EGL areas and REST^TG^ mice abnormal EGL areas (n=3). Scale bars in top panels apply to all panels within column. (D) Vimentin (E) Laminin (F) Gfap and (G) Blbp staining of P8 WT EGL areas and REST^TG^ mice abnormal EGL areas. Scale bars in top panels apply to all panels within a column.

Staining for the intermediate filament, Vimentin, also supported a disorganized migration of CGNPs in the cerebella of *REST^TG^* mice compared to WT animals (Fig. 1D. S1D) (52). IHC for Laminin identified perturbations in the integrity of the basement membrane around the cerebellum and within the fissures between folia in *REST^TG^* compared to WT mice (Fig. 1E, S1E). Further, glial cell projections (Gfap+ and Blbp+) were disorganized in areas with mis-migrating CGNPs (Figs. 1F, 1G, S1F-G). Gfap and Blbp were absent in meningeal pockets of infiltration (Figs. 1F-G, S1F-G). Thus, REST elevation in CGNPs resulted in cell autonomous and non-cell autonomous effects on cerebellar architecture.

### REST elevation in CGNPs promotes changes in the alternative transcriptome

To obtain a better understanding of the molecular mechanisms underlying the above processes, RNAseq analysis was performed using ex vivo cultured and CGNPs from brains of P8 WT and *REST^TG^* mice (n=3, each). *REST* FPKM was significantly but modestly elevated in *REST^TG^* CGNPs compared to WT cells (p=0.016) (Fig. 2A). qRT-PCR confirmed that this is likely attributable to the significantly higher expression of the h*REST* transgene in *REST^TG^* CGNPs compared to WT CGNPs (p<0.0001) (Fig. 2B). Endogenous mouse *Rest* expression was not significantly different between CGNPs from *REST^TG^* and WT mice (Fig. 2B).

**Figure 2.**
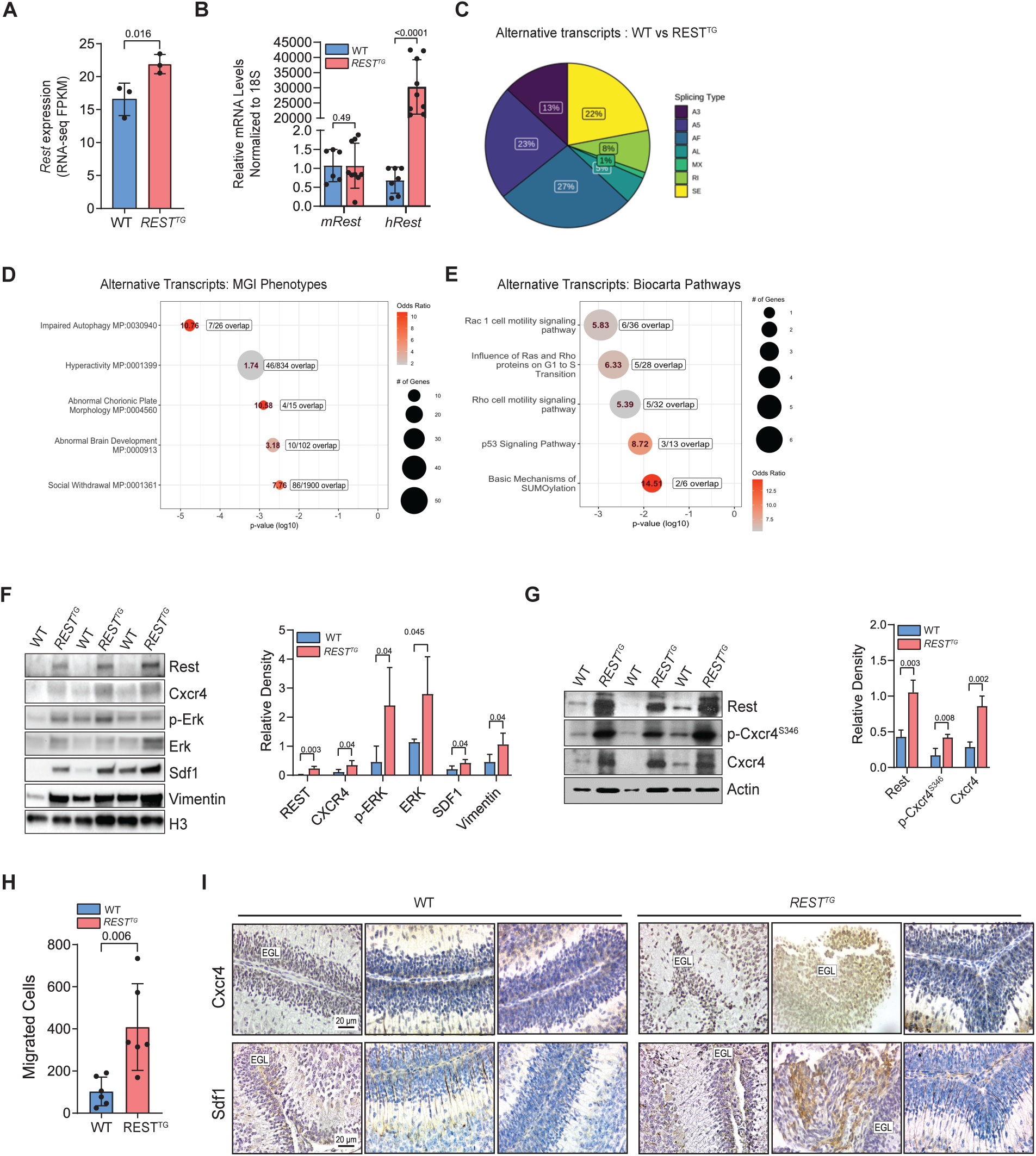
Mouse cGNPs (WT and REST^TG^) were analyzed by (A) RNA-seq (FPKM) to show human REST expression (n=3, each). P-values were calculated using unpaired student’s t-test. (B) qRT-PCR measurement of mouse Rest and human REST^TG^ mRNA expression in mouse (WT; n=7 and REST^TG^; n=9) cGNPs. (C) Pie chart of splicing events observed in REST^TG^ cGNPs; A3: alternative 3’ splice site (SS), A5: alternative 5’ SS, AF: alternative first exon, AL: alternative last exon, MX: mutually exclusive splicing, RI: retained intron, SE: skipped exon. (D and E) Top 5 enriched and nominally significant (p<0.05) MGI phenotypes (left) and molecular pathways (Biocarta, right) from differentially expressed alternative transcripts between WT and REST^TG^ cGNPs (yellow box highlight represents result with adjusted p-value <0.05). (F) Western blot to show differences in Cxcr4, Sdf1 and phospho-ERK, total ERK and actin loading control in proliferating WT (n=3) and REST^TG^ (n=3) cGNPs (left). Bar graph of densitometry quantification is shown on the right. (G) Western blot to demonstrate phosphorylation of CXCR4 C-terminal tail using S346 (mouse)/S339 (human) specific antibody (left) and densitometry quantification (right). (H) Number of WT and REST^TG^ cGNPs migrated to rSdf1 (170 ng/ml) as a chemoattractant at 24 hr (n=6). P values were calculated based on unpaired student’s t-test. (I) Representative 40x images of Cxcr4 and Sdf1 staining in sagittal brain sections of the WT (n=3) and REST^TG^ (n=3) cerebellar lamina.

Of the 727 genes with differential expression (p<0.05) between WT and *REST^TG^* CGNPs, 405 genes exhibited an absolute log2 fold change (log2FC) greater than 0.5 (226 upregulated) or lower than –0.05 (179 downregulated) (Fig. S2A). Pathway analysis showed an enrichment of processes relevant to altered brain and cerebellar development, cell adhesion, and β-arrestin dependent recruitment of kinases in GPCR signaling in *REST^TG^* over WT CGNPs (Figs. S2B-C).

A genome-wide Find Individual Motif Occurrences (FIMO) analysis was completed to identify potential direct REST targets using the canonical binding motif (MA0138-3) (Table S6). Of the 405 significantly altered genes, only 6 genes contained high-confidence (q<0.01) *RE1* binding motifs. This number was increased to 73 genes (28 genes ± 1 log2FC) using nominal statistical confidence (p<0.05). However, none of these candidates could explain the phenotypic changes with cell migration described in Fig. 1.

We also assessed the presence of alternative mRNA isoforms in our RNAseq dataset by performing a gene-level transcript analysis to identify genes exhibiting differential usage of alternative transcripts. This revealed a large number of genes (668) which contained isoform-specific alterations in expression in non-principal transcripts as determined by APPRIS (Table S7A). The differentially expressed alternative transcripts in *REST^TG^* CGNPs compared to WT cells were of the following types: alternative first (AF) exon (27%), alternative 5’ donor site (A5; 23%), skipped exon (SE; 22%), alternative 3’ donor site (A3; 13%), retained introns (RI; 8%), alternative last (AL) exon (5%) and mutually exclusive exons (MX; 1%) (Fig. 2C). Pathway analysis of genes containing differentially expressed alternative transcripts (n=668) further supported an enrichment of pathways relevant to brain development, autophagy, and Rho/Rac signaling and cell motility (Fig. 2D, 2E) (Table S7B). Thus, these results support a REST elevation-dependent alternative transcriptomic signature with potential for impact on cerebellar development and cell migration.

### REST elevation in CGNPs promotes Cxcr4-Sdf1 signaling hyperactivation and increases cell migration

To investigate the most phenotypically relevant pathway genes further, we evaluated the Abnormal Brain Development (MP:0000913) annotation which contained 42 genes with known relevance to either EGL expansion (MP: 0000873) or abnormal cerebellum EGL morphology (MP: 0000872) (Table S8A). Of these candidate genes, five (*Ptch1*, *Cxcr4*, *Egfr*, *Mdm4*, and Smarcb1) are implicated in MB genesis according to the cumulative evidence from Open Targets and an EGL-specific phenotype has been shown specifically for Ptch1 and Cxcr4 (Table S8B) (53-58).

Of note, Cxcr4-mediated migration towards Sdf-1/Cxcl12 enables CGNPs migration tangentially across the EGL prior to terminal migration and differentiation into the internal layer (17,18). Consistent with a potential role for Cxcr4/Sdf-1 signaling in REST-mediated mis-localization of CGNPs, Western blotting revealed a significant increase in Vimentin, Cxcr4, and Sdf1 levels (p=0.04, each) in *ex vivo* cultured *REST^TG^* CGNPs compared to WT cells (Fig. 2F). Interestingly, the increase in Cxcr4 protein levels (∼3.2-fold) was not accompanied by elevation in *Cxcr4* transcript expression, suggesting a post-translational mechanism of Cxcr4 activation (data not shown) (17,18). (Fig. 2G). This was confirmed by the >2-fold increase in Cxcr4 phosphorylated at serine 346 (pCxcr4^S346^) in *REST^TG^* compared to WT CGNPs (Fig. 2H). Cxcr4 phosphorylation at serine 346 is indicative of receptor activation, which also triggers downregulation of signaling by its endocytic internalization (33,59,60). However, sustained Cxcr4 signaling was observed in *REST^TG^* CGNPs which was functionally validated using transwell migration assays where a ∼3-fold increase in chemotactic response to rSDF1 was observed (Fig. 2H).

Cxcr4 and Sdf1 levels were also evaluated in the cerebella of WT and *REST^TG^* mice as well as in murine SHH (*Ptch^+/–^*) and REST-driven (*Ptch^+/–^/REST^TG^*) SHH MBs, because REST elevation in human SHH MBs is associated with increased risk for metastasis and poor survival (16). As shown in Fig. 2I, mis-localized *REST^TG^* CGNPs showed increased Cxcr4 staining and were found in the leptomeningeal space where higher Sdf1 staining was also noted. In contrast, Cxcr4 staining in the cerebella of WT mice was restricted to CGNPs in the outer EGL (Fig. 2I). In murine REST-driven Shh MBs, leptomeningeally disseminated tumors exhibited higher levels of Cxcr4, Sdf1, and Vimentin compared to *Ptch+/-* tumors (Fig. 3A-E). These findings support the potential relevance of REST-associated increase in this signaling axis to both cerebellar development and metastatic MB.

**Figure 3.**
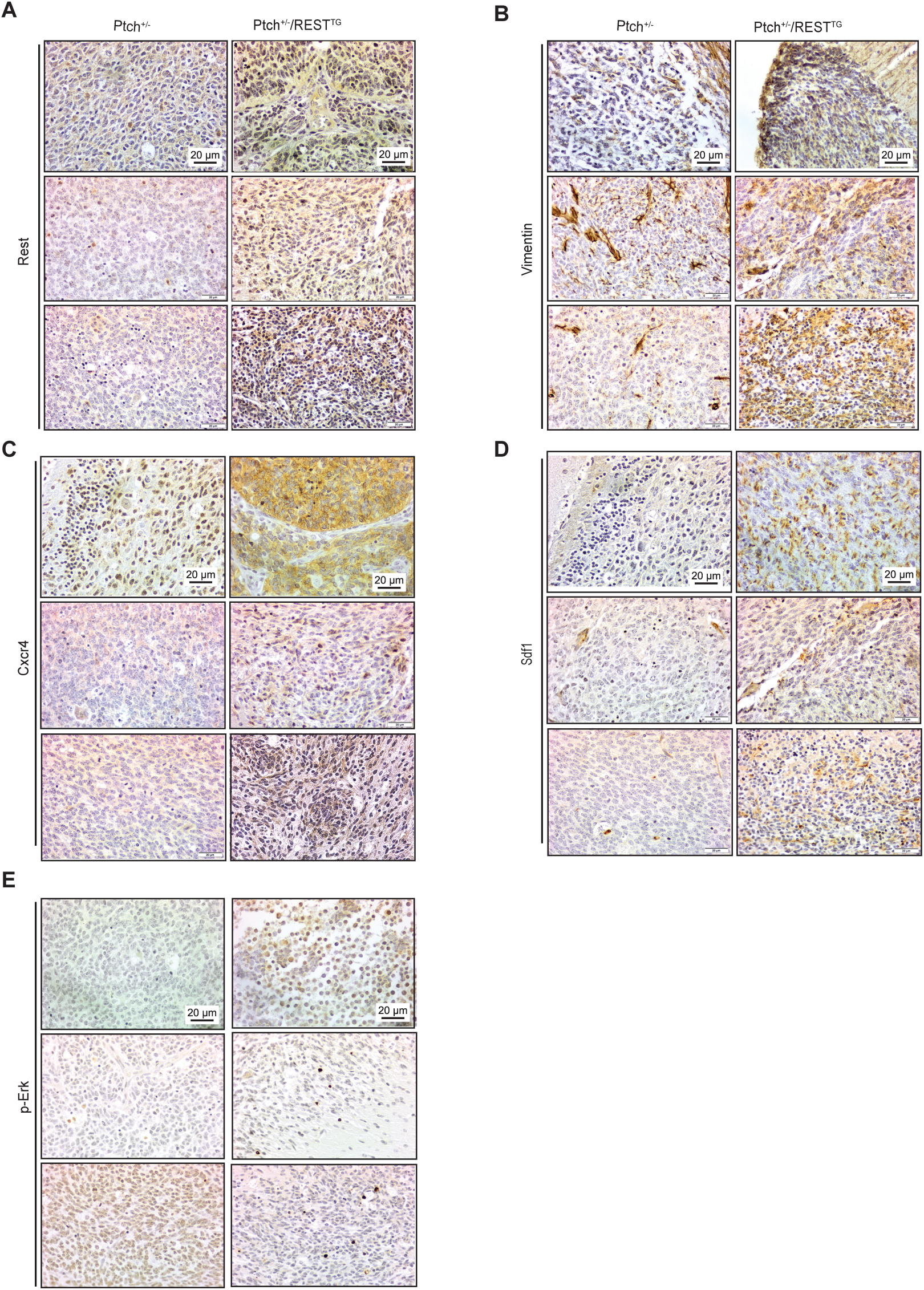
Representative immunohistochemistry panels to demonstrate changes in staining for (A) REST, (B) Vimentin, (C) Cxcr4, (D) SDF1, and (E) phosphorylated-ERK1/2 in tumors from Ptch^+/–^ and Ptch^+/–^/ REST^TG^ mice. Scale bars in top panels apply to all panels within column.

### REST elevation in CGNPs upregulates the expression of an alternative *Grk6* transcript

To define the mechanism by which REST elevation contributes to upregulated CXCR4 levels and activity, a list of known and predicted interactors and regulators of Cxcr4 were compiled using STRING (Table S9) (61). Of the 76 protein candidates that were identified, we focused on the G-protein coupled receptor kinase 6 (*Grk6*) initially because of the significant body of literature describing its role in the control of Cxcr4 stability and signaling activity in mice and humans (33,62,63).

Total *Grk6* levels were not significantly different between *REST^TG^* and WT CGNPs (Fig. 4A). At an isoform-level, 5 of the 7 Ensembl-annotated transcripts recognized for mouse *Grk6* were detected in the RNAseq data of WT and *REST^TG^* CGNPs (Fig. 4B-C). Of these 5 transcripts, only the FPKM of *Grk6*-207 was significantly increased in *REST^TG^* CGNPs versus WT cells (p=0.004) (Figs. 4B-C). Junction coverage analyses using the MISO algorithm applied to the RNAseq dataset confirmed detection of the 5 annotated *Grk6* isoforms in CGNPs from *REST^TG^* and WT mice (Figs. 4C, 4D). Again, only *Grk6*-207 – an isoform lacking exon 10a – demonstrated significantly increased junction coverage in *REST^TG^* CGNPs (average 25% junction coverage) versus WT (average 2% junction coverage) (Fig. 4D). These findings were confirmed by qPCR assays (Figs. 4E). Exon 10a encodes a part of the catalytic domain of Grk6 with relevance to Cxcr4 phosphorylation and binding (Fig. 4B, 4F). Other *Grk6* transcripts containing the same transcription start site and all the relevant domains were included in *Grk6*-203 (GenCode primary transcript) and *Grk6-*202. Averaged FPKM read coverage over each wildtype-like transcript (202 vs 203) did not exhibit differences between WT and *REST^TG^* CGNPs, and junction coverage tracks also did not reveal a major difference related to inclusion of the 3’ exon encoding the alternate 16 amino acids (residues 560-576) at the C-terminus of *Grk6*-202 (Figs. 4C, 4D).

**Figure 4.**
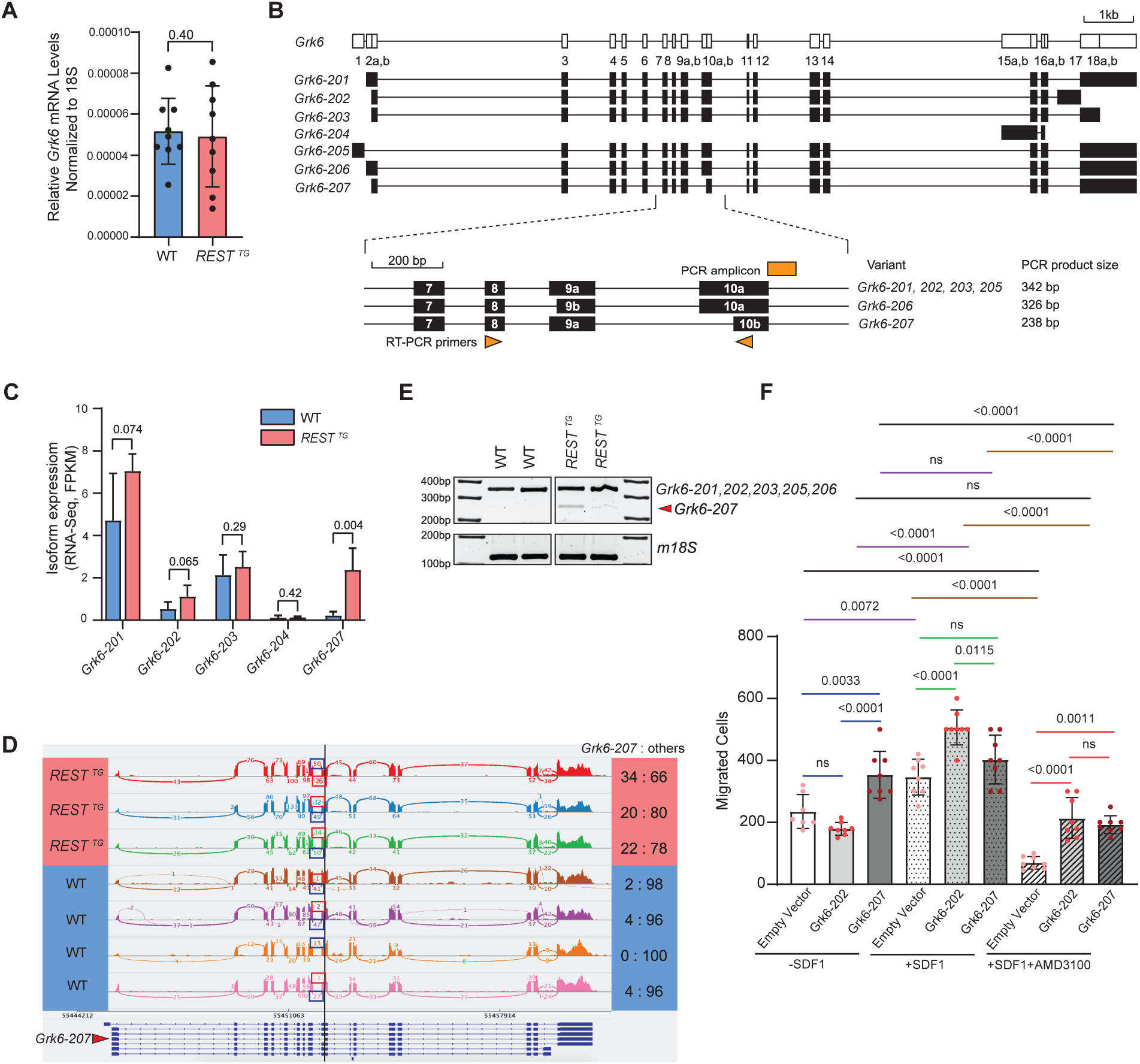
(A) qRT-PCR measurement of mouse *Grk6* in WT (n=9) and *REST^TG^* (n=9) cGNPs. (B) Schematic of mouse *Grk6* locus and alternative transcripts including location of primers to assess exon 10a exclusion. (C) Differential expression of *Grk6* isoforms -201, -202, -203, -204 and -207 measured by aligned RNA-seq read coverage averaged over the transcript bodies (FPKM) in mouse WT and *REST^TG^* cGNPs. P values were calculated based on unpaired student’s t-test. (D) Sashimi plots based on MISO algorithm representing alternative splicing in *Grk6.* P values were calculated based on unpaired student’s t-test. (E) qRT-PCR and agarose gel electrophoresis to confirm specific expression of a truncated 238bp band (arrow) corresponding to *Grk6* isoform - 207 in *REST^TG^*cGNPs compared to isoforms containing exon 10a (including *Grk6-201/203)*. 18S was used as a loading control. Primer locations are shown in B. (F) Transwell migration assay showing the number of WT and *REST^TG^* cGNPs migrating with and without the addition of Sdf1 (170 ng/ml) chemoattractant and Cxcr4 inhibitor (AMD3100) at 24 hr. Significance values were calculated based on multiple unpaired t-test.

Follow-up functional studies were done with WT CGNPs transduced with lentiviral vector expressing *Grk6*-202 or *Grk6*-207 to compare cell migration in the presence and absence of rSDF1 using transwell migration assays. Vector-transduced WT CGNPs or cells expressing *Grk6-*202 showed a comparable number of migrated cells in the absence of rSDF1, whereas CGNPs expressing *Grk6-*207 had a significantly higher number of migrated cells compared to vector-transduced or *Grk6-*202 expressing cells (p=0.0033 and <0.0001, respectively) (Fig. 4F). When compared to the above experiment conducted in the absence of rSDF1, the addition of rSDF1 caused a significant increase in the number of migrated WT CGNPs transduced with either empty vector or expressing *Grk6-*202 (p=0.0072 and <0.0001, respectively) but remained unchanged in *Grk6-*207 expressing cells (Fig. 4F). To test the requirement for Cxcr4-Sdf1 signaling for *Grk6-207*-induced increase in CGNP migration, the assay was performed in the presence of SDF1 and the CXCR4 inhibitor - AMD3100. As shown in Fig. 4F, in comparison to +rSDF1 alone conditions, cell migration could be significantly reduced in CGNPs transduced with vector alone or expressing *Grk6-202* or *Grk6-207* treated with AMD3100 (p=<0.0001 and <0.0001, respectively). These data indicate that the migration of CGNPs required rSDF1-dependent engagement and activation of Cxcr4 receptor, although its threshold for activation in the presence of *Grk6-207* was likely lower.

### Computational modeling predicts altered Grk6-Cxcr4-Arrb1 interactions due to REST-dependent exclusion of exon10a in *Grk6*

To understand how exon 10 exclusion in *Grk6* and expression of Grk6-207 may affect Cxcr4-Sdf1 signaling, we conducted an *in silico* structural biology analyses using AlphaFold3, an Artificial Intelligence (AI)-powered computational approach which predicts tertiary and quaternary protein structures using primary sequences (64,65). Primary sequences were obtained from NCBI and Uniprot for Cxcr4 (P70658), Grk6-202 (Q9EP84), Grk6-207 (A0A286YE10), and β-arrestin-1 (Arrb1; J3QNU6) and predictions were made with the presence of phosphorylated serine 346 (S346-p), ATP, and ADP.

Grk6-202 was predicted to interact with Arrb1 and Cxcr4 with a preference for Cxcr4-Arrb1 interaction especially when Grk6 is bound with ADP over ATP, similar to Grk6-203 (Fig. S3 A-B). The phosphorylated tail of Cxcr4 was accessible for interaction with the positively charged groove of Arrb1, as described in prior cryo-EM studies of phosphorylated Cxcr4 and Arrb1 (pTM = 0.46) (66). Thus, comparisons of Grk6-202 and Grk6-203 revealed highly similar profiles of constrained structural changes near the ATP-binding site which is relevant to phosphorylation of the C-terminal tail of Cxcr4 (Fig. S3C-D) (33,63) (67). We observed similar trends in less complicated predictions between only Cxcr4 and Grk6 isoforms 202/203 (pTM=0.67) and 207 (pTM = 0.54) (Fig. S4A-D).

Next, given that exon 10 exclusion in *Grk6*-207 has the potential to alter active site conformation and binding to ATP, AlphaFold3 was used to assess Grk6-207-Cxcr4-Arrb1 interactions (Fig. 5A). Interestingly, the interaction between Cxcr4 and Grk6-207 was predicted to be stronger than with Grk6-202, with the phosphorylated tail of Cxcr4 exhibiting an altered association near the ADP/ATP binding region and a potentially new predicted direct binding at arginine 341 (R341) (pTM = 0.43) (Fig. 5B-5E) Together, these results suggest that upregulated *Grk6-207* synthesis in *REST^TG^* CGNPs has the potential to increase the affinity of the gene product for phosphorylated Cxcr4 N-terminal residues and interfere with Arrb1-mediated Cxcr4 inactivation.

**Figure 5.**
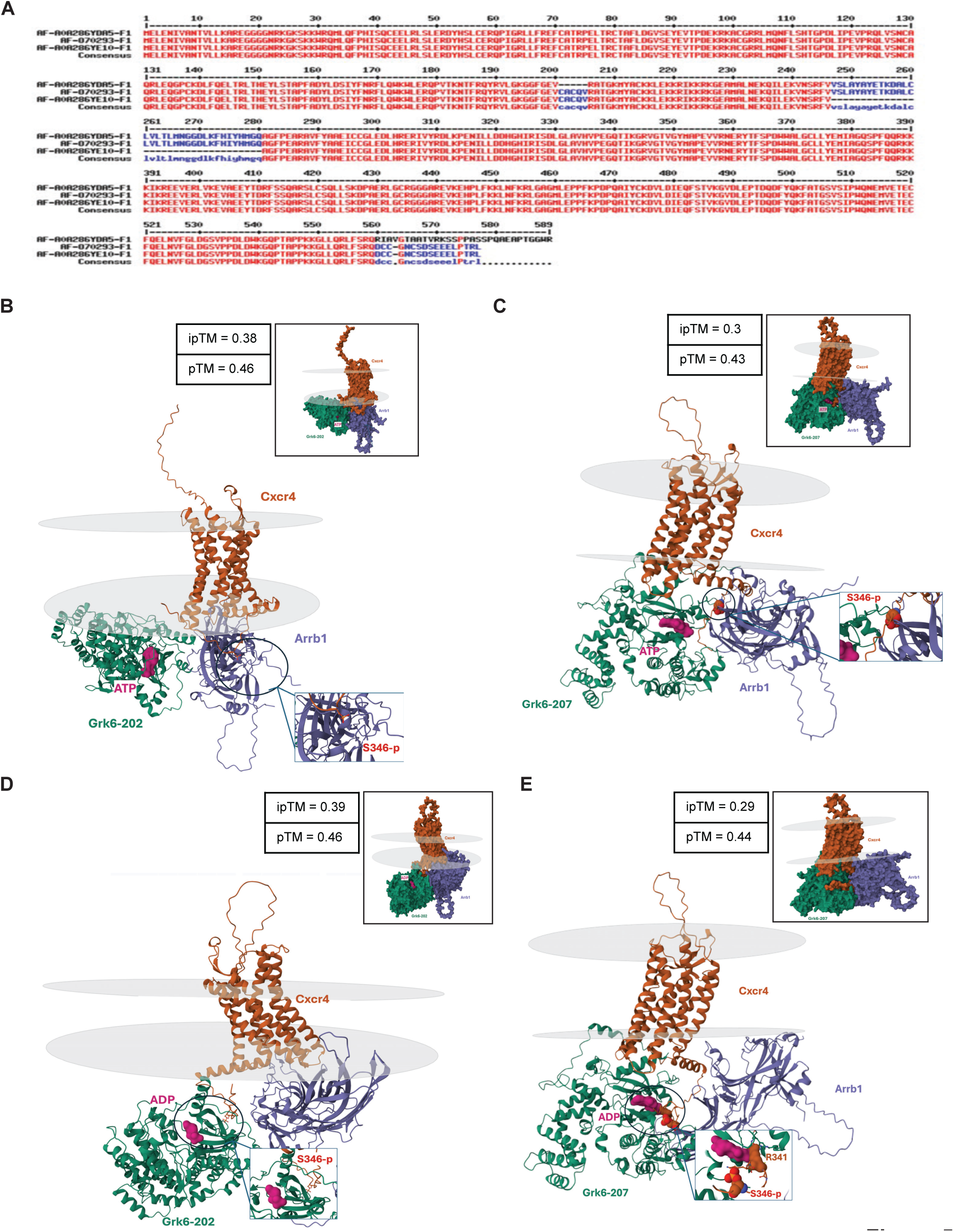
(A) Amino acid sequence comparison of gene products encoded by *Grk6*-201, 202 and 207 transcripts. Sequences lost by exon 10 deletion are shown by a dotted line. Stick/ribbon structures of the Alphafold3 structural predictions using amino acid sequences of mouse *Grk6-202* (WT) and Cxcr4 and Arrb1 in the presence of (B) ATP and (C) ADP. Stick/ribbon structures of the Alphafold3structural predictions with mouse Grk6-207 (with exon 10a exclusion) and Cxcr4 and Arrb1 in the presence of (D) ATP and (E) ADP. Grey discs represent borders of the plasma membrane bilayer. Surface representations of predictions are provided in upper border boxes. Zoomed view of the phosphorylated S346 is provided in bottom bordered boxes. In panel D, R341 is predicted to be bound to ADP.

### REST-elevation promotes enhanced chromatin accessibility at the *Grk6* locus

To understand how REST elevation contributes to exon 10a exclusion and *Grk6*-207 transcript synthesis, ATACseq was first performed to investigate potential changes in chromatin accessibility under conditions of REST elevation. An unexpected increase in chromatin accessibility at two regions at the *Grk6* locus was seen in *REST^TG^* (n=3) and WT (n=2) CGNPs. The first ATACseq peak was located near the 5’ end of the gene and appeared to correspond to the transcription start site (*TSS*) (Fig. 6A). Consistent with this, the 5’ ends of *Grk6* transcripts -202, -203, -205, -206, and - 207 also mapped to this region (Fig. 4B). Additionally, this *TSS*-proximal peak also exhibited an increase in the promoter specific histone H3 lysine (K)-4 trimethylation (me3) in *REST^TG^* CGNPs compared to WT cells by ChIPseq and confirmed by ChIP analysis (p<0.0001) (Figs. 6B-C). Surprisingly, while histone H3K27 acetylation (H3K27ac) was reduced at *Grk6 TSS* by ChIP and ChIPseq (p<0.0001), histone H3K9ac was increased in this region by both assays (p<0.0001) in *REST^TG^* CGNPs relative to WT cells (Figs. 6B-C).

**Figure 6.**
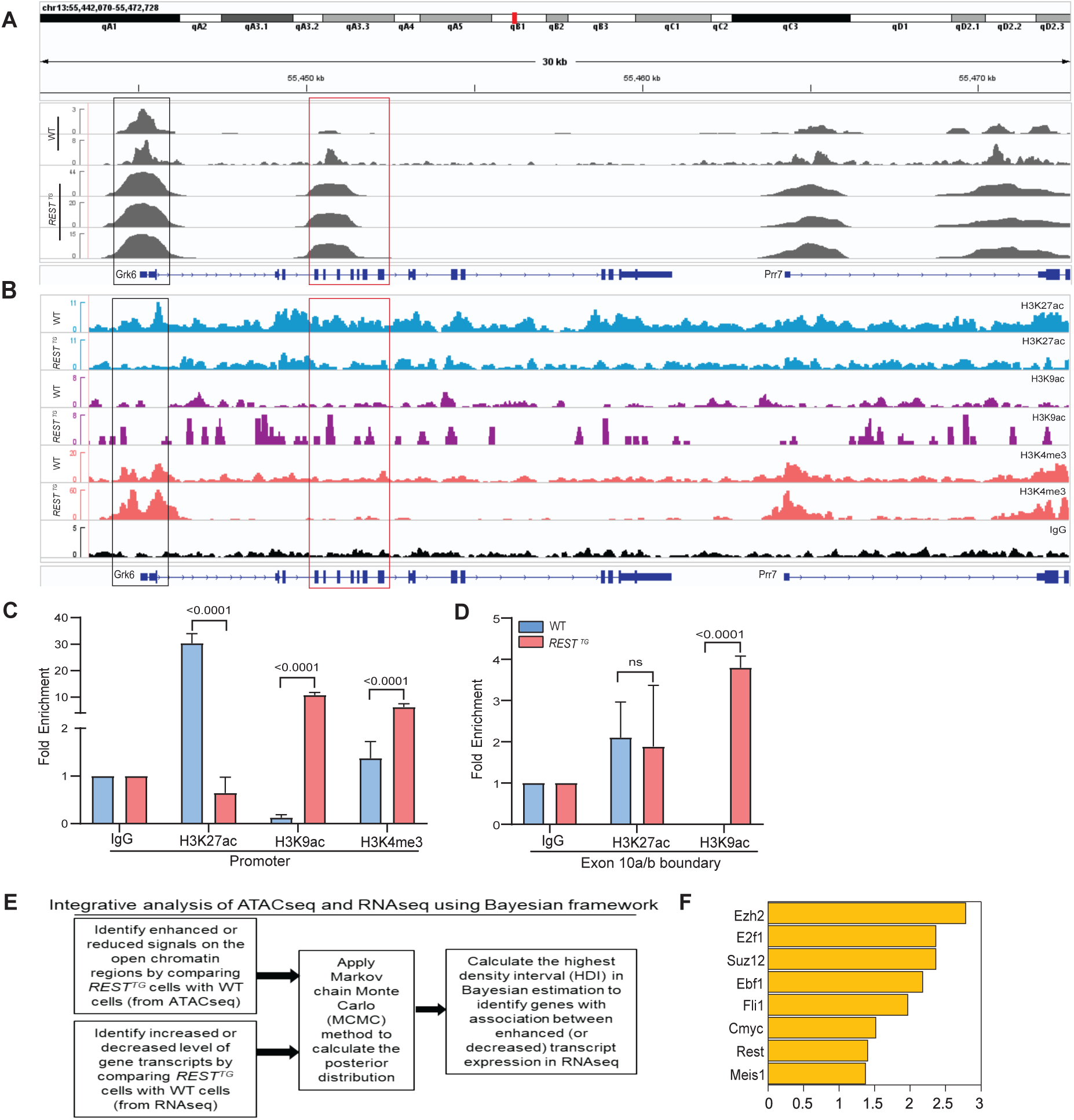
(A) Differences in chromatin accessibility at the *Grk6* locus on chromosome 13 in WT and *REST^TG^* CGNPs as measured by ATACseq analysis. (B) ChIPseq analysis was performed using pooled proliferating CGNPs from wildtype (WT) and *REST^TG^* mice (n=5, each) to assess deposition of the following histone modifications: H3K27ac, H3K9Ac, and H3K4me3 over a 30Kb region of chromosome 13, which includes the *Grk6* locus. IGV was used to visualize ATACseq and ChIPseq tracks. (C) ChIP-qPCR was done to provide changes in the enrichment of histone H3K4me3, H3K27ac, H3K9ac at the *Grk6* transcription start site and (D) in histone H3K27ac, H3K9ac at the exon 10a/b boundary of *Grk6*. P values were calculated using multiple comparisons in a Two-way Anova test. (E) Statistical framework for integrative Bayesian analysis. (F) Transcription Factor (TF) target enrichment analysis of 616 mouse TFs to demonstrate statistically significantly overlap between 427 genes with “coordinated epigenomics and transcriptome” patterns from panel E and target genes of 8 mouse TFs (Rest, E2f1, Ebf1, Fli1, Cmyc and Meis1) and Prc2 complex proteins (Ezh2 and Suz12).

The second ATACseq peak mapped to a region upstream of exon 10a of *Grk6*, which we also focused on given its potential relevance to exon 10a exclusion (Fig. 6A). Increased accessibility of chromatin 5’ of *Grk6-*exon 10a was not associated with a change in histone H3K27ac between *REST^TG^* and WT CGNPs (Figs. 6B-D). However, histone H3K9 acetylation showed a small increase in *REST^TG^* compared to WT CGNPs by ChIPseq and was further validated by ChIP, where an approximately 4-fold increase difference (p<0.0001) in the mark was observed between the two cell types (Figs. 6B-D). Thus, these data suggest that REST elevation in CGNPs promotes enhanced chromatin accessibility and a surprising preference for histone H3K9ac over histone H3K27ac at these regions, and that it results in an unexpected upregulation of *Grk6*-207 transcript expression in *REST^TG^* CGNPs. Positive and negative control regions for ATACseq and ChIPseq for histone H3K27ac, histone H3K9ac and histone H3K4me3 are shown in supplemental data (Fig. S5A-D).

### Integrated RNAseq/ATACseq analysis identifies a role for EZH2 in REST-dependent *Grk6* exon 10a exclusion

To identify candidate proteins with potential to bind chromatin under the ATACseq peaks 1 and 2, we adapted our previously published Bayesian-based statistical framework to perform an integrated analysis of the RNAseq and ATACseq datasets from *REST^TG^* and WT CGNPs (Fig. 6E) (40,41). This defined genes with statistically significant “coordinated epigenome and transcriptome” patterns based on stringent statistical criteria (probability for coordinated epigenome and transcriptome must be more than 99.9% in Bayesian estimation). We identified a total of 427 genes that met the above that passed Bayesian statistics threshold between *REST^TG^* and WT CGNPs, with 364 genes exhibiting differentially open ATACseq peaks at promoter regions and increased transcript expression, and 63 genes with diminished ATACseq peaks at promoter regions and decreased transcript expression (Table S10). Transcription factor (TF)-target enrichment analysis using published ChIPseq datasets of 616 mouse TFs collected from ENCODE and Cistrome datasets also showed statistically significantly overlap between 427 genes with “coordinated ATACseq and transcriptome” patterns from Fig. 6E and target genes of 8 mouse TFs (Rest, E2F1, Ebf1, Fli1, C-Myc and Meis1) and PRC2 complex proteins (Ezh2 and Suz12) (Fig. 6F and Fig. S5E).

Of these, Ezh2 was chosen as a candidate for further evaluation because of its known role in controlling CGNP differentiation and in MB genesis (68-70). Indeed, ChIPseq analyses revealed an increase in Ezh2 binding and a corresponding enhancement of histone H3K27me3 deposition (Table S11), and a concomitant reduction in histone H3K36me3 in the region of ATACseq peaks 1and 2 in *REST^TG^* CGNPs compared to WT cells (Fig. 7A). Positive and negative control regions for Ezh2, histone H3K27me3, and histone H3K36me3 are included in supplemental data (Fig. S6A, B).

**Figure 7.**
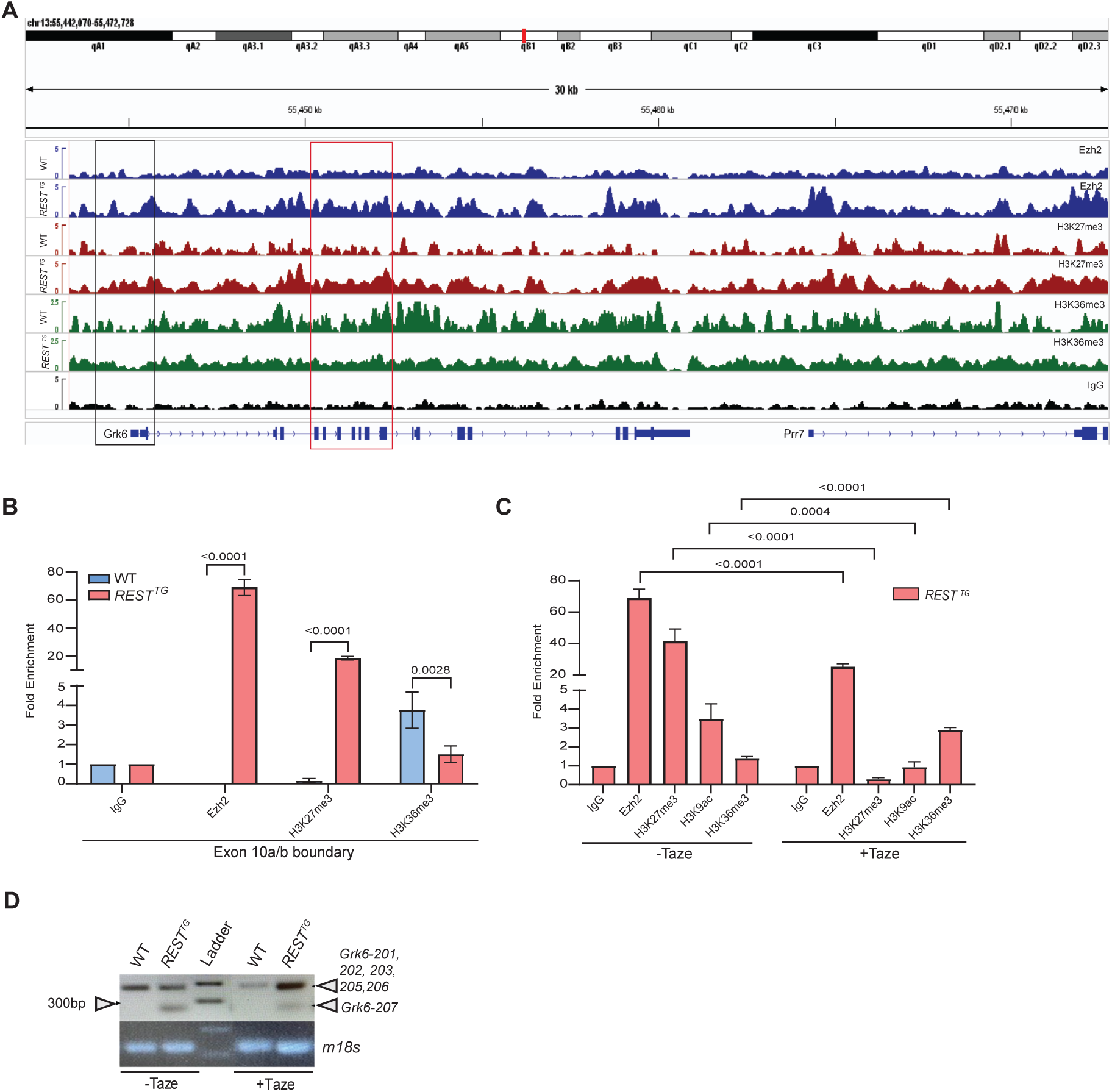
(A) ChIPseq was performed using proliferating CGNPs from wildtype (WT) and *REST^TG^*mice (n=8, each) to assess Ezh2 occupancy and to assess H3K27me3, and H3K36me3 (n=5, each). Tracks showing enrichment over a 30Kb region of chromosome 13, which includes the *Grk6* locus were visualized in IGV. (B) ChIP-qPCR of CGNPs from untreated WT (n=5) and *REST^TG^* (n=5) mice to assess fold-enrichment over IgG of Ezh2 and H3K27me3 at the *Grk6* exon 10a/10b boundary. (C) ChIP-qPCR to measure differences in enrichment of Ezh2, H3K27me3, H3K9ac, and H3K36me3 at *Grk6* REST^TG^ (n=5) CGNPs and in *REST^TG^* CGNPs (n=5) with or without treatment with Tazemetostat (taze). IgG was used for normalization. P values were calculated using multiple comparisons in a Two-way Anova test. (D) RT-PCR and agarose gel electrophoresis was used to evaluate differential amplification of a 238bp band (arrow) corresponding to *Grk6*-207 in *REST^TG^*CGNPs compared to a 342 bp band corresponding to exon 10a containing transcripts (including *Grk6-201/202/203/205)*. 18S was used as a loading control.

These findings were validated by ChIP analyses, where EZH2 occupancy and histone H3K27me3 deposition on chromatin at the *Grk6-*exon 10a-10b junction was significantly increased (p<0.0001, each) and histone H3K36me3 was significantly reduced (p=0.0028) in *REST^TG^* CGNPs compared to WT cells (Figs. 7B). Thus, REST elevation promotes an increase in Ezh2 occupancy and activity as well as a reduction in histone H3K36me3 at the Grk6 exon 10a-10b junction to impair transcription of exon 10a.

To establish the dependence of *Grk6* exon 10a exclusion on Ezh2 activity in *REST^TG^* CGNPs, cells were treated with the Ezh2 inhibitor, Tazemetostat (Taze) and ChIP analyses were performed. As shown in Fig. 7C, Taze treatment significantly reduced Ezh2 occupancy at the exon 10 site >3-fold (p<0.0001) and significantly decreased H3K27me3 (p<0.0001) relative to DMSO-treated controls. This was associated with a reduction in histone H3K9ac (p=0.0004) and restoration of histone H3K36me3 (p<0.0001) (Figs. 7C). Importantly, PCR analysis confirmed that Taze treatment reduced *Grk6-*207 expression in *REST^TG^* CGNPs indicating Rest-elevation driven increase in Ezh2 activity contributes to exon 10a exclusion *Grk6-*207 synthesis (Fig. 7D).

In summary, our studies implicate Rest elevation in driving significant changes in alternative transcript synthesis and attribute a role for Ezh2 and chromatin remodeling in Rest-dependent *Grk6* exon 10a exclusion/*Grk6*-207 synthesis. These findings have functional implications for the control of Cxcr4/Sdf1 signaling, CGNP migration, and postnatal cerebellar foliation.

## DISCUSSION

REST is a canonical transcriptional repressor heavily studied for its regulatory role in terminal neuronal differentiation in the brain cortex (71-73). The current study is one of the first to highlight its role in cerebellar development, specifically in the control of cerebellar foliation and CGNP migration. These findings align with published work from our group and others implicating *REST* elevation in leptomeningeal dissemination and metastasis of cerebellar SHH-MBs in mice and children with poor survival (8,15,16,71,72,74,75). Interestingly, we find Pax-6 positive CGNPs mis-localized in the cerebellar EGL near and within the leptomeningeal space of *REST^TG^* mice while the arrangement of other cell types and components of the extracellular matrix were also significantly perturbed. These data suggest the need for a better understanding of cell-cell communication and its contribution to foliation defects and loss of cerebellar symmetry in *REST^TG^* mice. Our findings may not be entirely surprising given that spatially and temporally constrained crosstalk between CGNPs, radial glia, meningeal cells, mature granule neurons, and interneurons is known to be important for normal cerebellar development (76,77). Nevertheless, our work provides a detailed characterization of a specific pathway through which *REST* elevation perturbs these coordinated processes: Cxcr4-Sdf1 signaling, an important axis in both cerebellar foliation and SHH-MB metastasis (17-20). Regulation of Cxcr4-Sdf1 signaling through post-translational modifications and chaperone-mediated endocytosis has been carefully dissected by other groups (33,59,60). However, our report is the first to implicate *REST* in the molecular regulation of the Cxcr4-Sdf1 axis, adding a potential epigenetic dimension to the control of this signaling pathway, which we will discuss later. It is also worth noting that future studies aiming to delineate the contributions of these findings to CNS tumors, in particular SHH-MB, will need to carefully consider the tumor microenvironment to assess the translational impact of these findings in the clinic (78,79).

REST has been largely studied as a repressor of transcriptional initiation and silencing of gene expression. Therefore, the significant change in the expression levels of alternative transcripts and the resultant increase in transcript diversity under conditions of *REST* elevation was somewhat surprising. A large fraction of these transcripts display alternative 5’ ends and alternative exon usage. Some of these also exhibit consensus REST-binding *RE1* sites in gene bodies suggesting that REST may be directly involved in the generation of transcript diversity. For example, *Nrxn1*, a brain-specific gene important in cerebellar and prefrontal cortex neuronal cell survival and implicated in neurological diseases (e.g., schizophrenia, Pitt Hopkins, autism) exhibits multiple intragenic *RE1* motifs and transcriptional diversity (80-82). However, the role of REST in driving alternative *Nrxn1* transcript generation and their involvement in the above neurological diseases in humans is currently unknown (83,84). Transcriptional diversity in the brain has been widely reported and given our findings here, a detailed evaluation of splice variants and specific isoform expression could potentially provide valuable insights into the etiology of neurological diseases (85-90). The continued identification and investigation of non-canonical regulators of alternative exon usage, such as the case we present here with REST, may be key to further decoding of relevant neuropathological mechanisms (91). Alternative transcripts are also shown to be significantly altered in human MBs, which is consistent with our findings in CGNPs, the cell of origin of SHH-MBs (92). These data support further analyses of REST and other potential regulators of exon usage in future studies.

The current study focused on *Grk6* because of its known role in the activation and desensitization of Cxcr4-Sdf1 signaling (33,62,63). Interestingly, we observed a REST-dependent increase in the synthesis of *Grk6-207*, a transcript containing an alternative transcriptional start site as well as alternative exons. Our data support a direct REST dependency of this synthesis given the presence of putative *RE1* sites at both the transcription start site and near the exon10a-10b region. Demonstration of REST occupancy of these loci would provide a direct role for REST in directing transcript diversity and is an area of ongoing effort in our group. However, we do show a role for REST-dependent changes in occupancy of other chromatin remodelers and histone modifications in exon 10a exclusion and synthesis of *Grk6-207*.

Our work is one of the first to implicate alternative transcripts of *Grk6* in CGNP migration. A combination of in silico computational modeling and functional studies was used to investigate the potential mechanisms underlying Grk6-207-mediated hyperactivation of Cxcr4-Sdf1 signaling and increased migration of CGNPs(33,93,94)￼. Structural analyses predict increased interaction between Grk6-207 and the C-terminal tail of Cxcr4 which may interfere with normal deactivating interactions of Cxcr4 with Arrb1. These computationally generated functional hypotheses are made possible by advances in deep learning models of protein structure (Alphafold3) trained on publicly available protein structures and their sequences(33,64,65)￼. An intriguing result of this analysis is that the interaction of Grk6-207 and the phosphorylated C-terminal tail of Cxcr4 is predicted to be stronger than that of this region with Grk6-202/203. In fact, in the presence of ADP, Grk6-207 is predicted to bind to a residue near S346 (R341) suggesting that loss of exon 10a may directly alter the docking and binding interactions in this region. A consequence of this increased affinity between Grk6 and Cxcr4 is that Arrb1 binding and subsequent inactivation is likely to be hindered(67,95)￼. Indeed, our functional migration assay supports this notion and additionally indicates that the activation threshold for Cxcr4 may be lower in CGNPs transduced with *Grk6-*207. Dedicated structural and biochemical studies are required to validate these computational estimations. It is worth noting that human orthologs of these proteins have very high homology, so these findings may also be conserved in humans and relevant to SHH-MBs, where CXCR4-SDF1 signaling is shown to drive metastasis(23)￼.

Congruent with REST’s function as a canonical repressor, increased REST activity is normally associated with chromatin compaction. However, our mechanistic studies uncovered a global increase in chromatin accessibility under conditions of *REST* transgene elevation in CGNPs. At the *Grk6* locus specifically, the ATACseq peak around the transcription start site is associated with increased deposition of histone H3K4me3, as well as an unexpected increase in histone H3K9ac, as opposed to histone H3K27ac. These data suggest a non-canonical transcriptional activating role for REST. Although speculative and needing validation, the increased chromatin accessibility seen in ATACseq data of *REST^TG^*mice combined with the accumulation of histone H3K9ac is suggestive of pause (and release) of the RNA polymerase II complex at promoters (96). RNA polymerase II pausing has been described at developmentally important genes and may be a reasonable regulatory event at *Grk6* given the importance of the Grk6-Cxcr4-Sdf1 axis in developmental control of cell migration in the cerebellum (97).

Nevertheless, this novel activating role for REST appears to be context dependent. In contrast to ATACseq peak near the transcription start site, the intragenic ATACseq peak is characterized by increased Ezh2 occupancy of chromatin and deposition of the repressive histone H3K27me3 mark and reduced histone H3K36me3, which is associated with exon 10a exclusion and *Grk6-207* transcript synthesis. This finding suggests that *REST* elevation-dependent upregulation of Ezh2 activity could potentially interfere with RNA processing. Whether this involves changes in splicing and or kinetics of RNA polymerase II (RNA PolII) during RNA elongation remains to be evaluated in future studies. Although histone H3K27me3 is not directly implicated in interaction with splice factors, it is known to indirectly influence this process by silencing the expression of splice factors (98). Other studies have shown that Ezh2 methylation of H3K27 creates a molecular roadblock to RNA Polymerase II-dependent elongation through the recruitment of histone H1 (99-101). Direct interaction of histone H3K36me3 with splice factors is known to modulate exon inclusion/exclusion (103,104). This interaction begets a fundamental question on what comes first at the *Grk6* exon 10a/10b boundary in *REST^TG^* CGNPs – a reduction in histone H3K36me3 or increase in histone H3K27me3, and how might REST direct these changes? Our future work looks to provide a better understanding of this process and to better understand which downstream target genes may be similarly impacted by these effects. Lastly, these effects are likely to be relevant to potential treatments in SHH-MB. Inhibition of Ezh2 methyltransferase activity, reducing H3K27me3, has been shown to stimulate MB cell differentiation and reduce tumor growth with potential as a treatment against SHH MB (102). The contributions of regulatory mechanisms and genetic targets contributing to these anti-tumor effects may be critical to understanding how Ezh2 treatment could be best utilized in the treatment of SHH MB patient segments.

In summary, we have uncovered a novel role for REST and chromatin remodeling in the control of alternative exon usage at *Grk6*, which has strong implications for regulation of CGNP migration, cerebellar foliation and development, and SHH MB (Figure 8).

**Figure 8.**
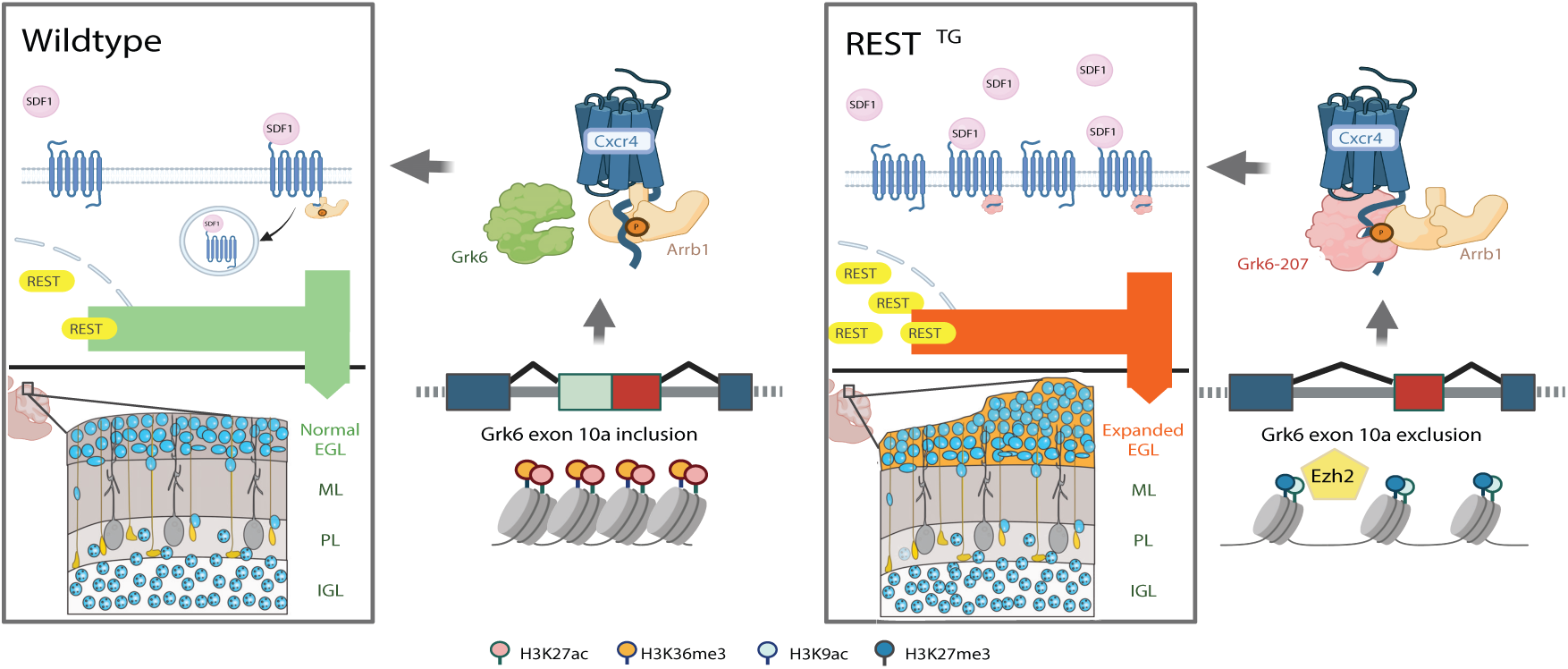
(Left panel) Physiological REST levels in CGNPs allow cell proliferation and tangential migration in the outer EGL regulated by Cxcr4-Sdf1 chemokine signaling. Developmentally controlled decline in *REST* expression reduces proliferation, initiates neurogenesis and downregulates Cxcr4-Sdf1 signaling activity to promote a switch from EGL-centric tangential to radial migration towards the IGL. This decline in Cxcr4-Sdf1 levels is caused by a Grk6 gene product encoded by the *Grk6-202* transcript, which features exons 10a and 10b, permits recruitment of Arrb1 to the Grk6-Cxcr4 complex and inactivation of chemokine signaling. (Right panel) In contrast, in *REST^TG^* mice, constitutive *REST* expression in CGNPs results in their hyperproliferation, blockade of neurogenesis and aberrant maintenance of tangential migration in the EGL to drive excessive accumulation of CGNPs in the EGL. This REST-dependent phenotypic change in Cxcr4-Sdf1 signaling and cell migration stems from abnormal Ezh2 occupancy and histone H3K27me3 deposition at the exon10a-10b locus in *Grk6*, and synthesis of an alternative transcript - *Grk6*-207 that lacks exon10a. *Grk6*-207 is proposed to encode a gene product with potential to interfere with Cxcr4 binding to Arrb1 and downregulation of Cxcr4-Sdf1 signaling. Created with BioRender.

## AUTHOR CONTRIBUTIONS

KC, JS: Conception, experimental design and execution of work, writing and manuscript edits, funding support; KC, LG, XH: bioinformatic analysis and execution of work, writing and manuscript edits; TD, ASi, JBA, ARH, YY: experimental design and execution of work and manuscript edits; ASh: reagent generation; LX: bioinformatics, oversight of work, manuscript edits, funding support; VG: Conception, experimental design, oversight of work, writing and manuscript edits, funding support.

## SUPPLEMENTARY DATA

Supplementary Data are available at NAR online.

## CONFLICT OF INTEREST

No competing interests.

## FUNDING

This work was supported by grants to KC from the National Center for Advancing Translational Sciences-National Institutes of Health (NIH) (TL1TR000369 and UL1TR000371); to LX from the Rally Foundation, the Cancer Prevention Research Institutes of Texas (CPRIT-RP180805) and NIH (R21CA259771 and P30CA142543); to VG from the Cancer Prevention Research Institutes of Texas (CPRIT-RP150301) and NIH (R01NS079715 and R03NS077021).

## DATA AVAILABILITY

RNA-seq, ATAC-seq and ChIPseq datasets will be uploaded to NCBI GEO omnibus (Accession numbers: TBD).

## Notes

### Competing Interest Statement

The authors have declared no competing interest.

