## Supplementary material for "REST elevation-dependent chromatin remodeling and alternative Grk6 transcript synthesis hyperactivates Cxcr4-Sdf1 signaling in cerebellar granule cell progenitors": Table S1, Table S2, Table S3, Table S4, Table S5

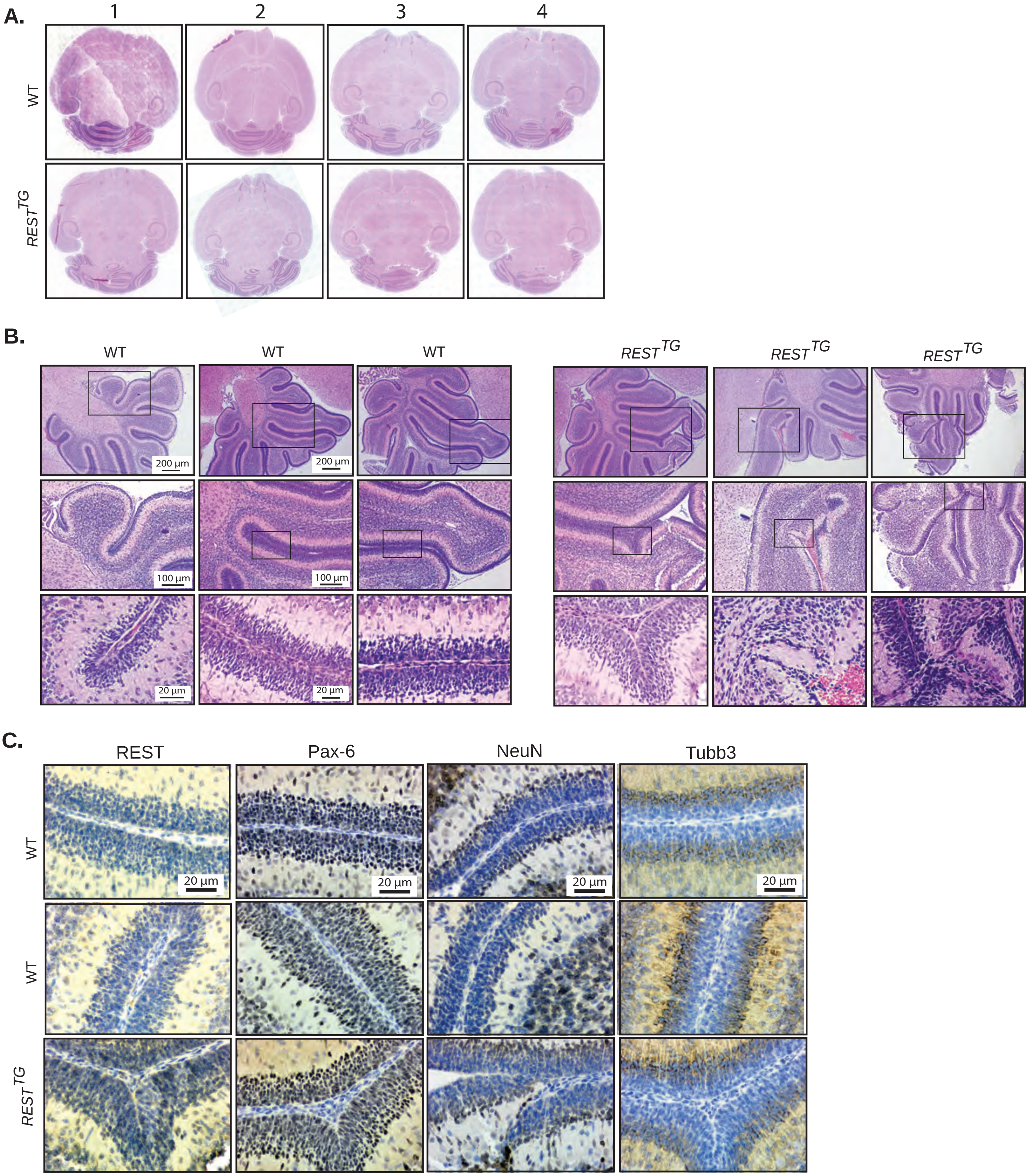

A.
1
2
3
4
WT
REST^TG
B.
WT
WT
WT
REST^TG
REST^TG
REST^TG
200 µm
200 µm
100 µm
100 µm
20 µm
20 µm
C.
REST
Pax-6
NeuN
Tubb3
WT
WT
REST^TG
20 µm
20 µm
20 µm
20 µm


Supp. Fig. 1

D.

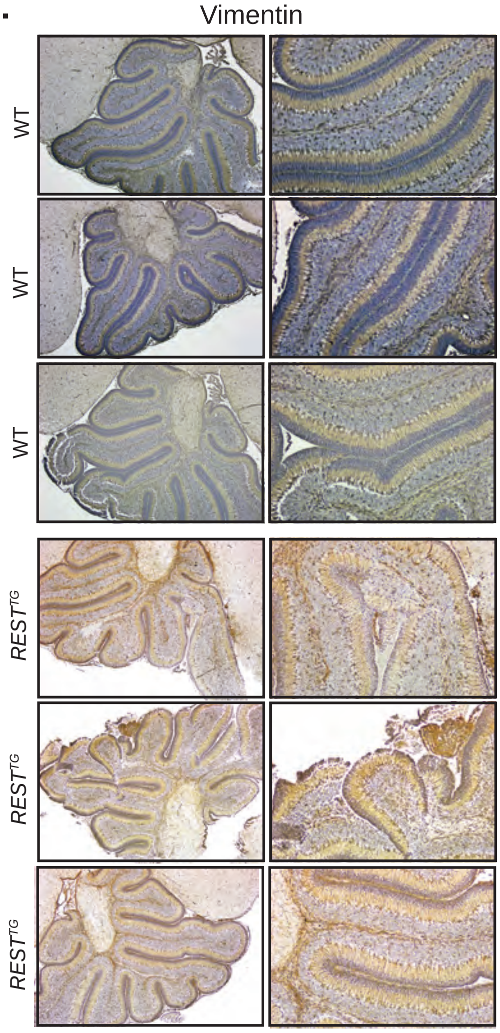


E.

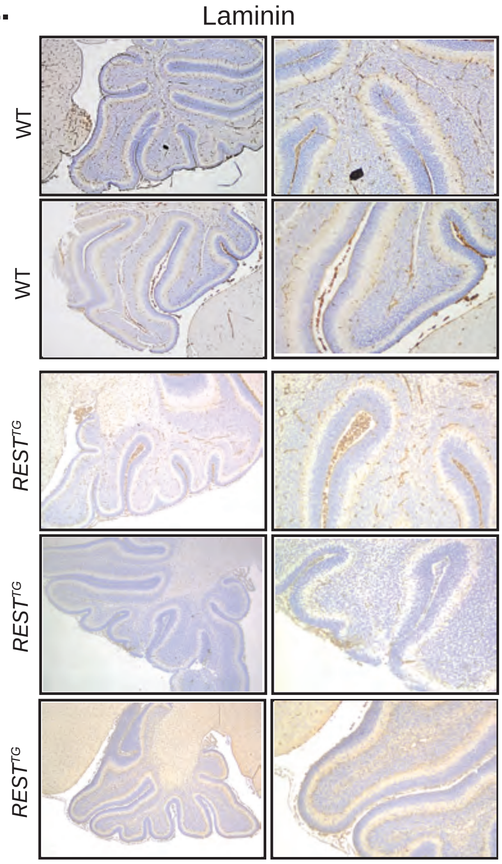


F.

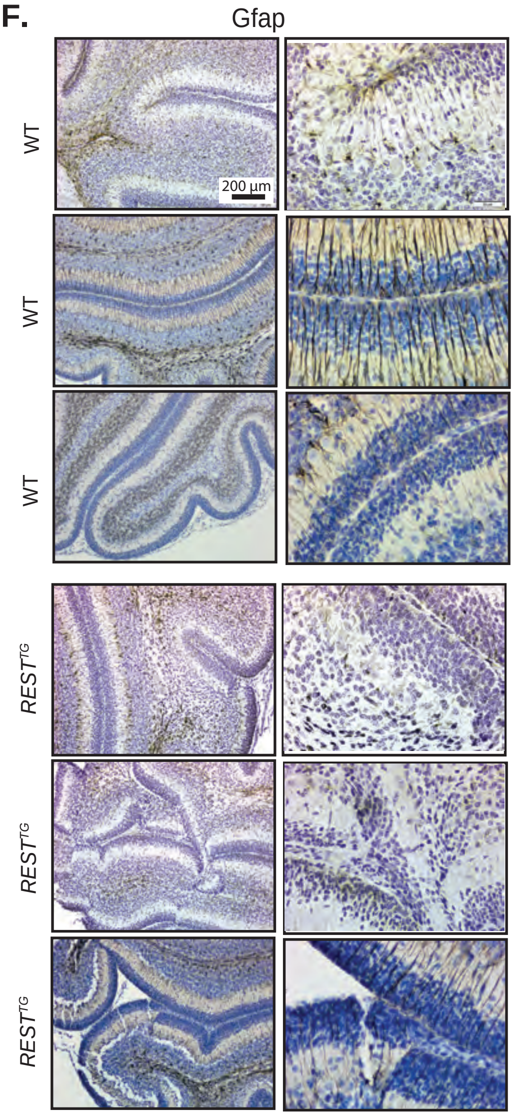


G.

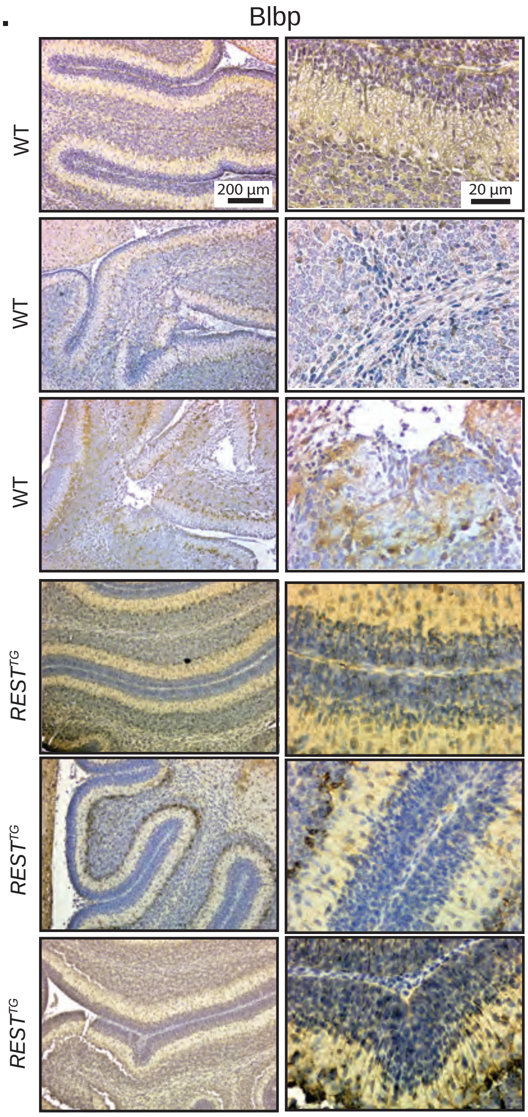


Supp. Fig. 1

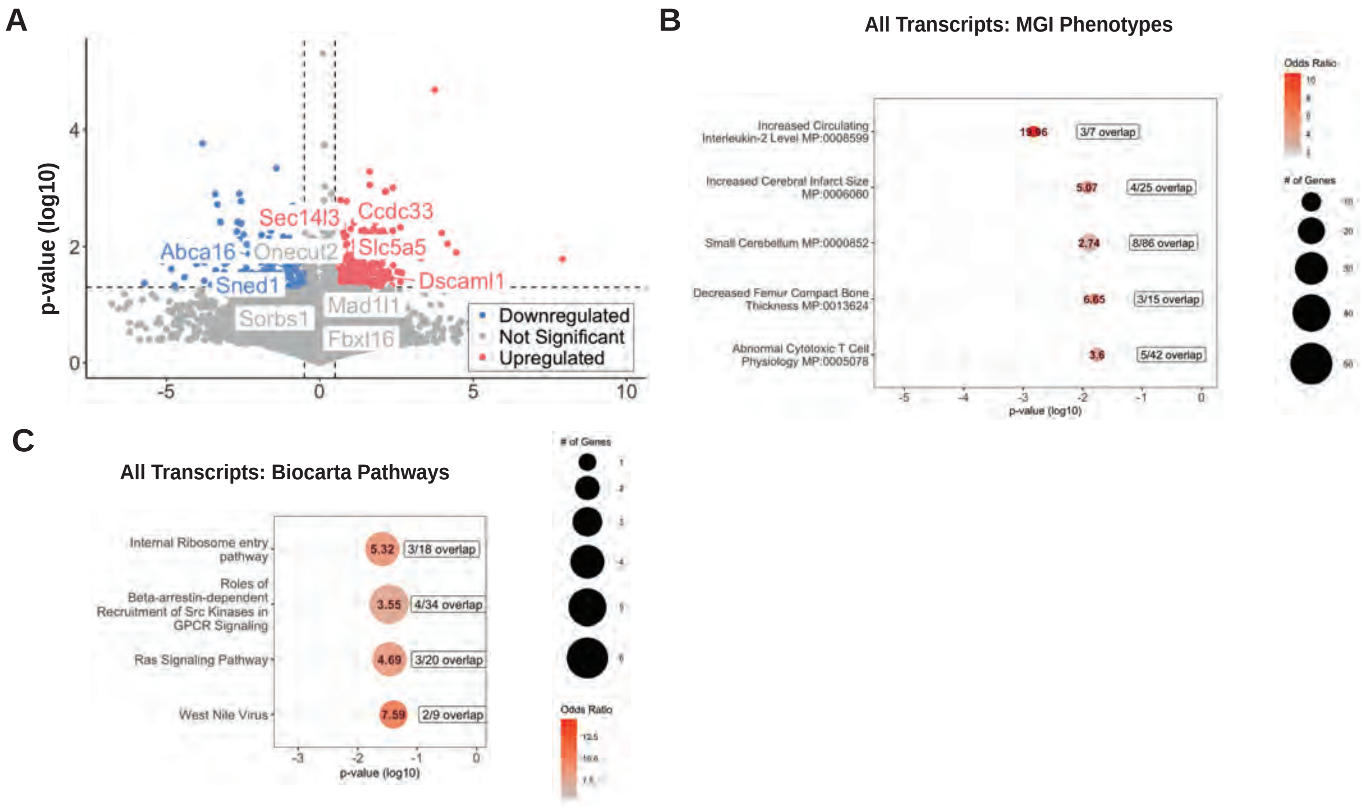
A
p-value (log10)
Sec14l3
Ccdc33
Abca16
Onecut2
Slc5a5
Sned1
Dscaml1
Sorbs1
Mad1l1
Fbxl16
Downregulated
Not Significant
Upregulated
B
All Transcripts: MGI Phenotypes
Increased Circulating Interleukin-2 Level MP:0008599
3/7 overlap
Increased Cerebral Infarct Size MP:0006080
5.07
4/25 overlap
Small Cerebellum MP:0000852
2.74
8/86 overlap
Decreased Femur Compact Bone Thickness MP:0013624
6.65
3/15 overlap
Abnormal Cytotoxic T Cell Physiology MP:0005078
3.6
5/42 overlap
p-value (log10)
Odds Ratio
### of Genes
C
All Transcripts: Biocarta Pathways
Internal Ribosome entry pathway
5.32
3/18 overlap
Roles of Beta-arrestin-dependent Recruitment of Src Kinases in GPCR Signaling
3.55
4/34 overlap
Ras Signaling Pathway
4.69
3/20 overlap
West Nile Virus
7.59
2/9 overlap
p-value (log10)
### of Genes
Odds Ratio

Supp. Fig. 2

Supplemental Figure 2A. RNA-sequencing pathway analysis.

Genes with putative RE-1 motifs ($p<0.05$)

| No. | Gene names | RNAseq FC | No. | Gene names | RNAseq FC |
|---|---|---|---|---|---|
| 1 | Abhd3 | -0.6995336 | 43 | Pex5l | -0.6319134 |
| 2 | Actg2 | 1.41578462 | 44 | Pla2g7 | -1.2099169 |
| 3 | Adam18 | -1.1801238 | 45 | Plxnb1 | -0.5217478 |
| 4 | Adgrb1 | -0.580416 | 46 | Psg27 | 1.12132482 |
| 5 | AI467606 | 0.75599525 | 47 | Rgl1 | 0.53987986 |
| 6 | Aldh3a1 | -2.2369835 | 48 | Rims3 | 0.58440633 |
| 7 | Arhgap22 | -1.1576724 | 49 | Rtn4rl2 | 0.77305452 |
| 8 | Asic1 | 0.65879247 | 50 | Scrt1 | -0.8519393 |
| 9 | Bmp7 | 0.53336413 | 51 | Selenbp1 | 0.96091995 |
| 10 | Btc | -1.3249227 | 52 | Serpinb6d | 1.08920012 |
| 11 | Cabp5 | -0.7418395 | 53 | Sh3rf3 | 0.56187138 |
| 12 | Car12 | -0.8338592 | 54 | Slc22a18 | -1.1387966 |
| 13 | Casp8 | -0.5014259 | 55 | Slc43a2 | -0.9742032 |
| 14 | Ccdc148 | -0.9121784 | 56 | Slc8a1 | -0.5959781 |
| 15 | Ceacam19 | 1.72496176 | 57 | Slco6d1 | -1.4307103 |
| 16 | Col8a1 | -0.9984369 | 58 | Spaca3 | -1.3905089 |
| 17 | D630003M2Rlk | -0.719174 | 59 | Spats2l | 0.75242876 |
| 18 | Efna2 | 0.51038 | 60 | Spdl1 | 0.51862748 |
| 19 | Ehbp1l1 | -1.1900377 | 61 | Srd5a2 | -1.1834461 |
| 20 | Eln | 0.57239964 | 62 | Taar8a | -2.2733348 |
| 21 | Enkur | -0.8510465 | 63 | Tnfsf14 | 1.08443682 |
| 22 | Fcgbp | 0.6683151 | 64 | Tppp2 | 1.74020861 |
| 23 | Fev | 1.69746166 | 65 | Trabd2b | -1.0176281 |
| 24 | Fgf11 | 0.59104378 | 66 | Trim30c | -2.1764693 |
| 25 | Fgf18 | 0.81632677 | 67 | Trpm8 | 0.69151487 |
| 26 | Frmd6 | 0.52545867 | 68 | Unc93a2 | 0.56692521 |
| 27 | Galnt6 | -1.0587295 | 69 | Vav1 | -0.71402 |
| 28 | Gem | -1.0587295 | 70 | Vmn2r107 | -2.0006126 |
| 29 | Hnf1a | 0.83678178 | 71 | Wnt7b | -0.5071865 |
| 30 | Hsf5 | 0.65288483 | 72 | Zdhhc11 | 1.0196227 |
| 31 | Ireb2 | -0.619702 | | | |
| 32 | Islr | -1.0817079 | | | |
| 33 | Itgb4 | 0.56790881 | | | |
| 34 | Mov10l1 | 0.75920618 | | | |
| 35 | Mrap2 | -0.7363836 | | | |
| 36 | Mrgpra1 | -2.7067668 | | | |
| 37 | Mycbpap | 0.97284589 | | | |
| 38 | Myo16 | -0.5179541 | | | |
| 39 | Nmur1 | -1.1027255 | | | |
| 40 | Nt5c1b | 1.29378181 | | | |
| 41 | Oc90 | 1.04724261 | | | |
| 42 | Pde6b | 0.69100674 | | | |

Supplemental Figure 2B. RNA-sequencing pathway analysis.

| No. | Genes with putative RE-1 motifs (p<0.05) & significant DE (p<0.05) | RNAseq FC |
|---|---|---|
| 1 | Actg2 | 1.41578462 |
| 2 | Adam18 | -1.1801238 |
| 3 | Aldh3a1 | -2.2369835 |
| 4 | Arhgap22 | -1.1576724 |
| 5 | Btc | -1.3249227 |
| 6 | Ceacam19 | 1.72496176 |
| 7 | Ehbp1l1 | -1.1900377 |
| 8 | Fev | 1.69746166 |
| 9 | Galnt6 | -1.0587295 |
| 10 | Islr | -1.0817079 |
| 11 | Mrgpra1 | -2.7067668 |
| 12 | Nmur1 | -1.1027255 |
| 13 | Nt5c1b | 1.29378181 |
| 14 | Oc90 | 1.04724261 |
| 15 | Pla2g7 | -1.2099169 |
| 16 | Psg27 | 1.12132482 |
| 17 | Serpinb6d | 1.08920012 |
| 18 | Slc22a18 | -1.1387966 |
| 19 | Slco6d1 | -1.4307103 |
| 20 | Spaca3 | -1.3905089 |
| 21 | Srd5a2 | -1.1834461 |
| 22 | Taar8a | -2.2733348 |
| 23 | Tnfsf14 | 1.08443682 |
| 24 | Tppp2 | 1.74020861 |
| 25 | Trabd2b | -1.0176281 |
| 26 | Trim30c | -2.1764693 |
| 27 | Vmn2r107 | -2.0006126 |
| 28 | Zdhhc11 | 1.0196227 |

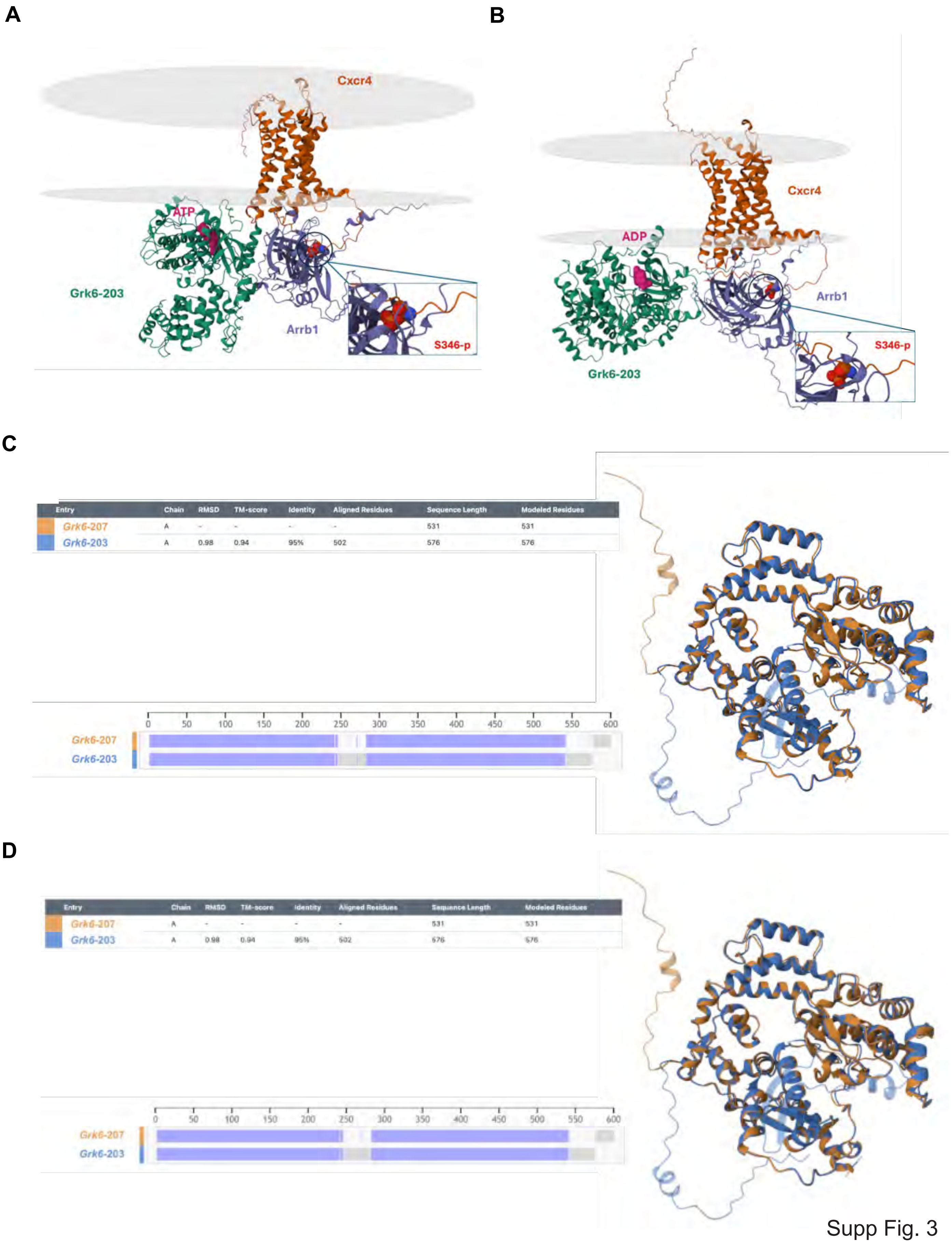
A
Cxcr4
ATP
Grk6-203
Arrb1
S346-p
B
Cxcr4
ADP
Grk6-203
Arrb1
S346-p
C
Entry Chain RMSD TM-score Identity Aligned Residues Sequence Length Modeled Residues
Grk6-207 A - - - - 531 531
Grk6-203 A 0.98 0.94 95% 502 576 576
0 50 100 150 200 250 300 350 400 450 500 550 600
Grk6-207
Grk6-203
D
Entry Chain RMSD TM-score Identity Aligned Residues Sequence Length Modeled Residues
Grk6-207 A - - - - 531 531
Grk6-203 A 0.98 0.94 95% 502 576 576
0 50 100 150 200 250 300 350 400 450 500 550 600
Grk6-207
Grk6-203

Supp Fig. 3

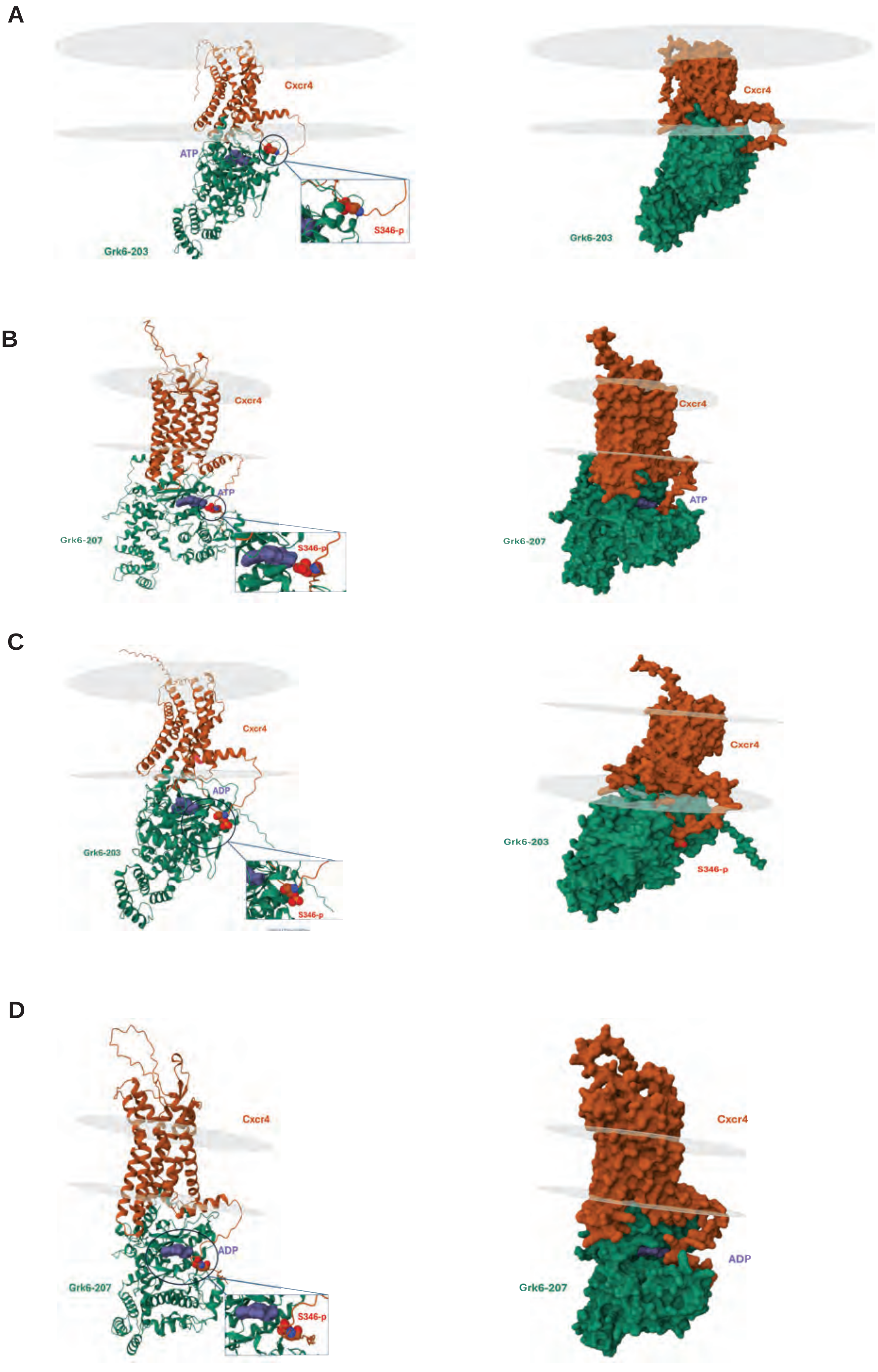
A
Cxcr4
ATP
S346-p
Grk6-203
Cxcr4
Grk6-203
B
Cxcr4
ATP
Grk6-207
S346-p
Cxcr4
ATP
Grk6-207
C
Cxcr4
ADP
Grk6-203
S346-p
Cxcr4
Grk6-203
S346-p
D
Cxcr4
ADP
Grk6-207
S346-p
Cxcr4
ADP
Grk6-207

Supp. Fig. 4

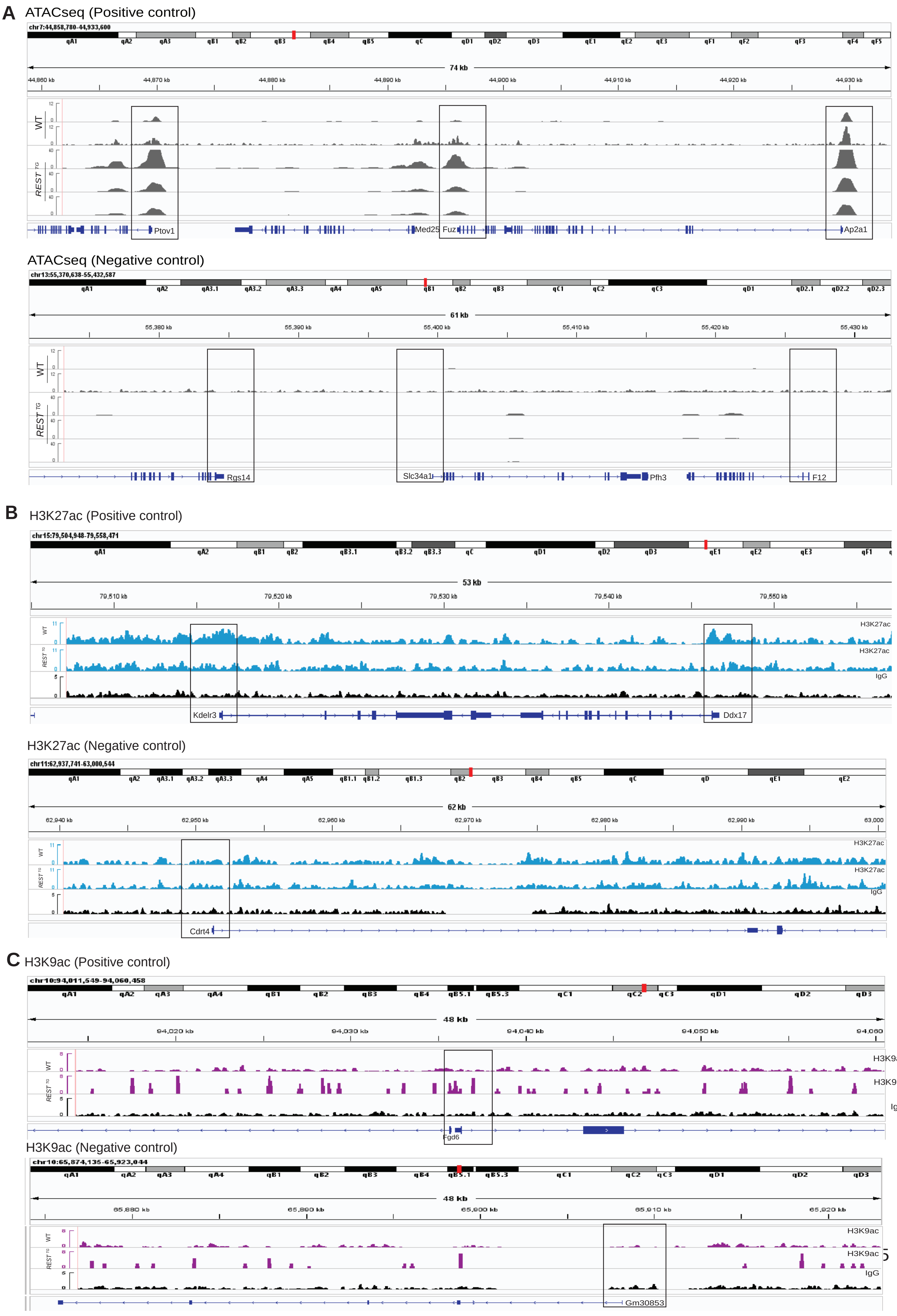
A
ATACseq (Positive control)
chr7:44,858,780-44,933,600
74 kb
WT
REST TG
Ptov1
Med25
Fuz
Ap2a1
ATACseq (Negative control)
chr13:55,370,638-55,432,587
61 kb
Rgs14
Slc34a1
Pfh3
F12
B
H3K27ac (Positive control)
chr15:79,504,948-79,558,471
53 kb
H3K27ac
IgG
Kdelr3
Ddx17
H3K27ac (Negative control)
chr11:62,937,741-63,000,544
62 kb
Cdrt4
C
H3K9ac (Positive control)
chr10:94,011,549-94,060,458
48 kb
H3K9ac
Fgd6
H3K9ac (Negative control)
chr10:65,874,135-65,923,044
48 kb
Gm30853

**D** H3K4me3 (Positive control)

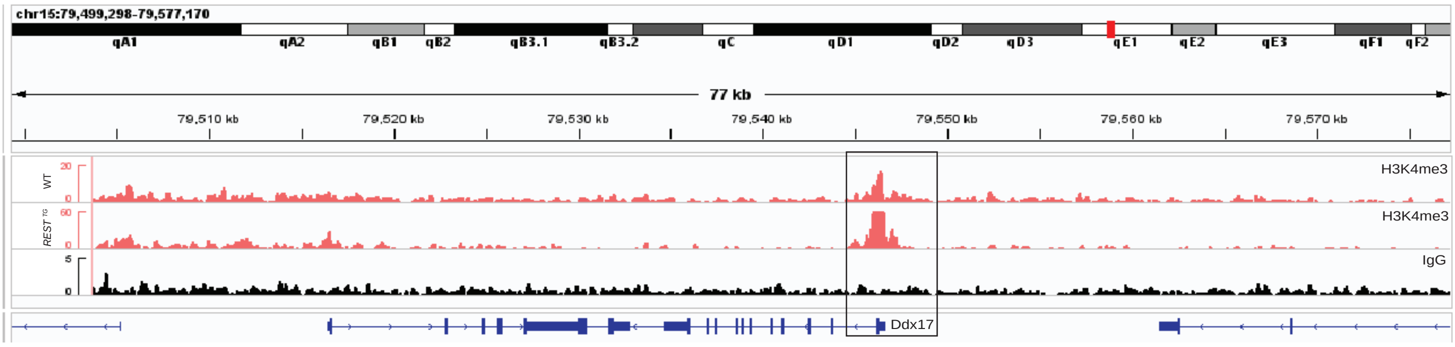


H3K4me3 (Negative control)

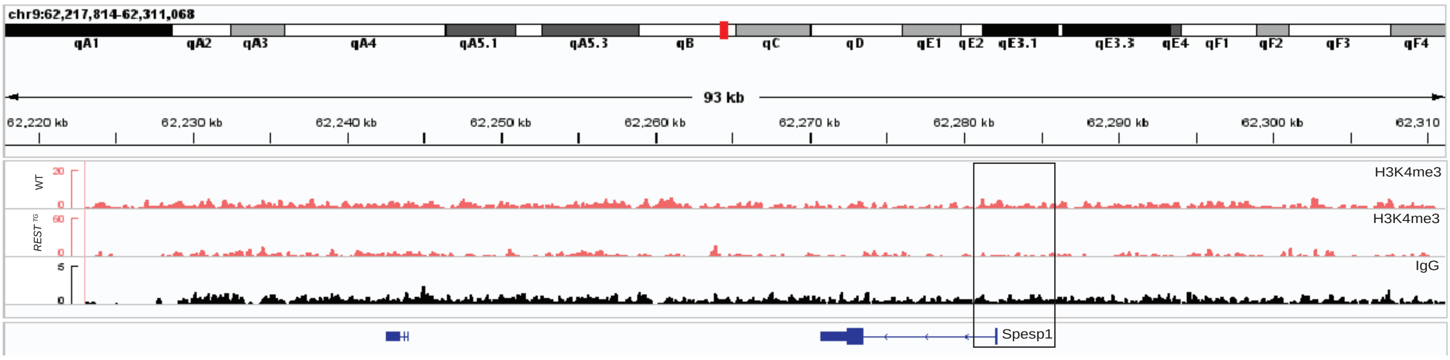


**E** Ezh2 ChIPseq target genes (mouse)

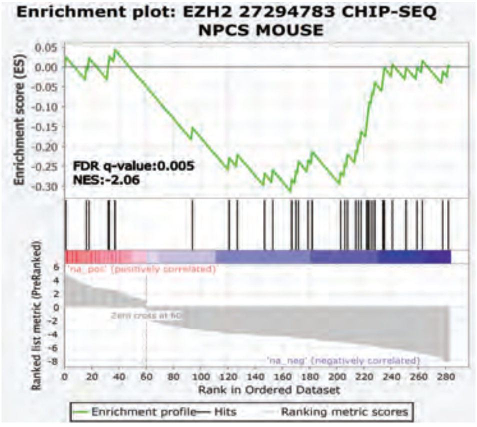

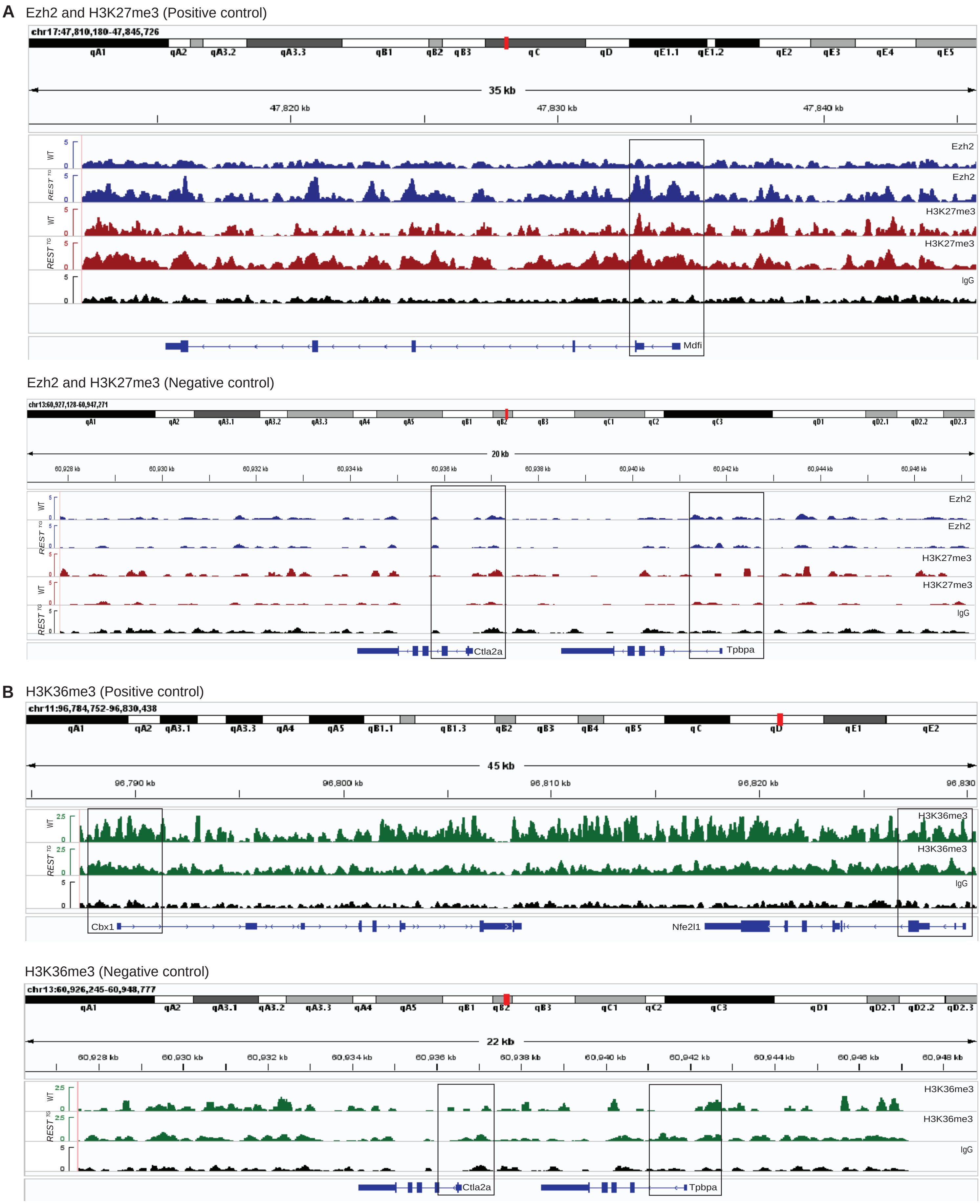

A
Ezh2 and H3K27me3 (Positive control)
chr17:47,810,180-47,845,726
35 kb
47,820 kb
47,830 kb
47,840 kb
WT
REST TG
Ezh2
H3K27me3
IgG
Mdfi
Ezh2 and H3K27me3 (Negative control)
chr13:60,927,128-60,947,271
20 kb
Ezh2
H3K27me3
IgG
Ctla2a
Tpbpa
B
H3K36me3 (Positive control)
chr11:96,784,752-96,830,438
45 kb
96,790 kb
96,800 kb
96,810 kb
96,820 kb
96,830
H3K36me3
IgG
Cbx1
Nfe2l1
H3K36me3 (Negative control)
chr13:60,926,245-60,948,777
22 kb
60,928 kb
60,930 kb
60,932 kb
60,934 kb
60,936 kb
60,938 kb
60,940 kb
60,942 kb
60,944 kb
60,946 kb
60,948 kb
H3K36me3
IgG
Ctla2a
Tpbpa


Supp. Fig. 6

Supplementary figure legends:

**Figure S1. REST overexpression disrupts normal cytoarchitecture in the postnatal cerebellum. (A)** H&E of additional samples of horizontal brain slices from Wildtype (WT) and *REST*$^{TG}$ mice used for symmetry analysis. **(B)** H&E staining of sagittal sections of cerebella from WT and *REST*$^{TG}$ mice in two additional *REST*$^{TG}$ lineages. Boxes in upper panels correspond to the frame of the next row. Scale bars in left panels apply to all panels within row. IHC for **(C)** REST, Pax6, NeuN, Tubb3, **(D)** Vimentin, **(E)** Laminin **(F)** Gfap, and **(E)** Blbp in sagittal sections of the cerebella of additional WT and *REST*$^{TG}$ mice. Scale bars in top panels apply to all panels within column.

**Figure S2. Differential gene expression and pathway analysis of RNAseq data of wildtype and *REST*$^{TG}$ CGNPs.**

**(A)** Volcano plot of gene expression between WT and *REST*$^{TG}$ cGNPs with highlighted genes representing those with significant differential expression $\geq \pm 1$ Log2FC. **(B)** Top 5 enriched and nominally significant ($p<0.05$) mouse genome informatics (MGI) phenotypes (left) and **(C)** all significantly enriched molecular pathways (Biocarta; right) for all nominally significant genes with differential expression ($p<0.05$.

**Figure S3. Comparison of Alphafold3 stick/ribbon structural predictions for Grk6-203 and Grk6-207.**

Stick/ribbon structures of the Alphafold3 structural predictions using amino acid sequences of Grk6-203 (WT) and canonical mouse Cxcr4 and Arrb1 in the presence of **(A)** ATP and **(B)** ADP. Grey discs represent borders of the plasma membrane bilayer. Surface representations of predictions are provided in upper border boxes. Zoomed view of the phosphorylated S346 is provided in bottom bordered boxes. **(C) and (D)** Variation in primary sequences of Grk6-203 and Grk6-207 are shown. Adjacent ribbon/stick figures are superimposed structures of Grk6-203 and Grk6-207 showing the differences.

**Figure S4. Comparison of Alphafold3 stick/ribbon structural predictions for Grk6-203 and Grk6-207 interaction with Cxcr4 in the presence ATP or ADP.**

Stick/ribbon structures and surface plots of the Alphafold3 structural predictions using amino acid sequences of Grk6-202 (WT) and canonical mouse Cxcr4 in the presence of ATP **(A)** and **(C)** and in the presence of ADP **(B)** and **(D)** ADP. Grey discs represent borders of the plasma membrane bilayer. Surface representations of predictions are provided in upper border boxes. Zoomed view of the phosphorylated S346 is provided in bottom bordered boxes.

**Figure S5. Positive and negative controls for ATACseq and ChIPseq analyses in WT and *REST$^{TG}$* CGNPs.**

**(A)** Differences in chromatin accessibility at positive (*Ptov1, Fuz* and *Ap2a1*) and negative (*Rgs14 and Slc34a*) control gene loci in WT and *REST$^{TG}$* CGNPs was measured by ATACseq. **(B)** Enrichment of H3K27ac at positive (*Kdelr3* and *Ddx17*) and absence of enrichment in negative controls genes (*Cdrl4*) are shown. **(C)** Enrichment of H3K9ac at positive (*Fgd6*) and its absence in negative controls genes (*Gm30853*) are shown. **(D)** Enrichment of H3K4me3 at positive (*Ddx17*) and its absence in negative controls genes (*Spesp1*) are shown. **(E)** GSEA analysis to identify enrichment of Ezh2 ChIPseq target genes among 427 coordinated genes.

**Figure S6. Positive and negative controls for ChIPseq analysis for EZH2 occupancy and enrichment of H3K27me3 and H3K36me3 in WT and *REST$^{TG}$* CGNPs.**

**(A)** Enrichment of Ezh2 and H3K27me3 at positive (*Mdfi*) and its absence in negative controls genes (*Ctla2a* and *Tpbpa*) are shown. **(B)** Enrichment of H3K36me3 at positive (*Cbx1* and *Nfe2l1*) and absence of enrichment in negative controls genes (*Ctla2a* and *Tpbpa*) are shown.

**Table S1**

**Antibodies - IHC**

| **Antibody** | **Catalog Number** | **Dilution** | **Company** |
|---|---|---|---|
| REST | 00141 | 1:50 | Bethyl Laboratories, Montgomery TX, USA |
| Vimentin | ab92547 | 1:200 | Abcam, Cambridge, UK |
| Laminin | ab11575 | 1:200 | Abcam, Cambridge, UK |
| Tubb3 | 801201 | 1:250 | Biolegend, San Diego CA, USA |
| NeuN | ab13938 | 1:200 | Abcam, , Cambridge, UK |
| Gfap | 3670 | 1:50 | Cell Signaling Technology, Danvers, MA, USA |
| Sdf1 | MAB350 | 1:100 | R&D Systems, Minneapolis MN, USA |
| Pax-6 | PRB-278P | 1:200 | Biolegend, San Diego CA, USA |
| Blbp | ab32423 | 1:200 | Abcam, Cambridge, UK |

**Table S2**

**Antibodies - Western**

| **Antibody** | **Catalog number** | **Dilution** | **Company** |
|---|---|---|---|
| REST | 07-579 | 1:1000 | Millipore, Burlington, MA, USA |
| ERK1/2 | 9102 | 1:1000 | Cell Signaling Technology, Danvers, MA, USA |
| p-Erk | 9101 | 1:5000 | Cell Signaling Technology, Danvers, MA, USA |

| | | | |
|---|---|---|---|
| p-CXCR4 | 59028S | 1:1000 | Cell Signaling Technology, Danvers, MA, USA |
| p-CXCR4 | ab74012 | 1:1000 | Abcam, Cambridge, UK |
| CXCR4 | ab124824 | 1:2000 | Abcam, Cambridge, UK |
| Sdf1 | MAB350 | 1:500 | R&D Systems, Minneapolis MN, USA |
| Vimentin | ab92547 | 1:1000 | Abcam, Cambridge, UK |
| Actin | ab7810 | 1:50000 | Abcam, Cambridge, UK |

**Table S3**

**qRT-PCR primers mouse CGNPs (WT vs REST$^{TG}$)**

| **Primer Name** | **Primer Sequence** |
|---|---|
| mGrk6 IG For | CACCTTCAGGCAGTACCGAG |
| mGrk6 IG Rev | CACAGCAGATCTCAGCAGCA |
| m18S F1617 | GCAATTATTCCCCATGAACG |
| m18S R1739 | GGCCTCACTAAACCATCCAA |
| mGrk6 For 1 | TGAGATGGTGGAGACCGAGT |
| mGrk6 Rev 1 | CTGCAGTTCCCACAGCAATC |
| mGrk6 For 2 | TTTGCCGACTACCTCGACAG |
| mGrk6 Rev 2 | GTACATCTTGCCTGTTGCCC |

**Table S4**

**Antibodies - ChIP**

| **Antibody** | **Catalog No.** | **Dilution** | **Company** |
|---|---|---|---|
| H3K9Ac | 9649 | 1:200 | Cell Signaling Technology, Danvers, MA, USA |
| H3K27me3 | 9733 | 1:200 | Cell Signaling Technology, Danvers, MA, USA |
| H3K36me3 | 4909 | 1:200 | Cell Signaling Technology, Danvers, MA, USA |
| H3K27ac | 8173 | 1:200 | Cell Signaling Technology, Danvers, MA, USA |
| Ezh2 | 5246 | 1:200 | Cell Signaling Technology, Danvers, MA, USA |

**Table S5**

**qPCR primers - ChIP**

| **Primer name** | **Primer Sequence** |
|---|---|
| mGrk6 TSS1-F | CAGAGGCTCGGCCCTAATTG |
| mGrk6 TSS1-R | CGATTTCTGGTTGGACTGCG |
| RE1 mGrk6 F | ACGCATTGTGTACAGGTAGGG |
| RE1 mGrk6 R | GAGCTGCCACTCTCAAACCT |
